# Deciphering Cell-types and Gene Signatures Associated with Disease Activity in Rheumatoid Arthritis using Single Cell RNA-sequencing

**DOI:** 10.1101/2023.10.05.560352

**Authors:** M. Binvignat, B. Y. Miao, C. Wibrand, M.M. Yang, D. Rychkov, E. Flynn, J. Nititham, W. Tamaki, U. Khan, A. Carvidi, M. Krueger, E. Niemi, Y. Sun, G. Fragiadakis, J. Sellam, E. Mariotti-Ferrandiz, D. Klatzmann, A. Gross, J. Ye, A. J. Butte, L.A Criswell, M. Nakamura, M. Sirota

## Abstract

**Objective:** Single cell profiling of synovial tissue has previously identified gene signatures associated with rheumatoid arthritis (RA) pathophysiology, but synovial tissue is difficult to obtain. This study leverages single cell sequencing of peripheral blood mononuclear cells (PBMCs) from patients with RA and matched healthy controls to identify disease relevant cell subsets and cell type specific signatures of disease.

**Methods:** Single-cell RNA sequencing (scRNAseq) was performed on peripheral blood mononuclear cells (PBMCs) from 18 RA patients and 18 matched controls, accounting for age, gender, race, and ethnicity). Samples were processed using standard CellRanger and Scanpy pipelines, pseudobulk differential gene expression analysis was performed using DESeq2, and cell-cell communication analysis using CellChat.

**Results:** We identified 18 distinct PBMC subsets, including a novel IFITM3+ monocyte subset. CD4+ T effector memory cells were increased in patients with moderate to high disease activity (DAS28-CRP ≥ 3.2), while non-classical monocytes were decreased in patients with low disease activity or remission (DAS28-CRP < 3.2). Differential gene expression analysis identified RA-associated genes in IFITM3+ and non-classical monocyte subsets, and downregulation of pro-inflammatory genes in the Vδ subset. Additionally, we identified gene signatures associated with disease activity, characterized by upregulation of pro-inflammatory genes *TNF, JUN, EGR1, IFIT2, MAFB, G0S2*, and downregulation of *HLA-DQB1, HLA-DRB5, TNFSF13B*. Notably, cell-cell communication analysis revealed upregulation of immune-associated signaling pathways, including VISTA, in patients with RA.

**Conclusions:** We provide a novel single-cell transcriptomics dataset of PBMCs from patients with RA, and identify insights into the systemic cellular and molecular mechanisms underlying RA disease activity.

## INTRODUCTION

Rheumatoid arthritis (RA) is a systemic autoimmune disease characterized by chronic inflammation and joint destruction (1). While the prevalence and disease burden vary considerably between geographic regions and populations (2), RA impacts approximately 1.3 million adults in the United States, representing 0.6 to 1% of the country’s population (3,4). RA is a debilitating condition and a major socio-economic burden with a prevalence of work disability around 35% (5). Effective RA management necessitates early diagnosis, a treat-to-target approach, and the attainment of remission or low disease activity (6). Achieving optimal therapeutic success remains the main challenge in RA as only 16 % of patients reach sustained remission or low disease activity (7,8). This has been particularly underscored by the recent recommendations from the European Alliance of Associations for Rheumatology (EULAR), particularly concerning the management of difficult-to-treat RA patients (9).

The understanding of cellular and molecular mechanisms underlying disease activity has garnered significant attention. Notably, specific cell subsets, such as synovial tissue macrophages, have been associated with both remission and disease activity (10). Additionally, several synovial molecular and pathobiological markers have shown promise in predicting treatment response (11–13). The emergence of bulk transcriptomic data has further revealed that alterations in synovial and blood transcriptomic profiles were closely associated with disease activity and flares (14–16). In the pursuit of comprehensive insights, single-cell RNA sequencing (scRNA-seq) emerges as a powerful tool to simultaneously profile cell subset compositions and cell type-specific transcriptional states, enabling a deeper understanding of mechanisms associated with non-remission. Several studies have utilized single-cell resolution to investigate RA, although the majority of research has focused on synovial tissue and none specifically studied disease activity (17–19).

Recent cross-tissue meta-analyses of transcriptome data, encompassing samples from both human and murine models, have uncovered genes associated with disease activity, and underscored a divergence between synovium and peripheral blood profiles (20,21). Consequently, it is crucial to recognize that markers identified in the synovium cannot be directly extrapolated to those found in peripheral blood. The accessibility of peripheral blood, compared to the more invasive nature of synovial sampling, emphasizes its practical advantage for both research and potential clinical applications. An additional challenge in studying RA disease activity is the inherent heterogeneity of the condition and the potential influence of demographic factors, such as gender, age, ethnicity and race, on disease activity(22–25). A critical aspect of addressing this challenge involves promoting the establishment of more standardized and diverse cohorts, enabling a better exploration of specific cell subsets and biomarkers that contribute to disease activity.

In this study, we describe a comprehensive analysis of disease activity using scRNA-seq of peripheral blood mononuclear cells (PBMC) in a diverse cohort of RA patients, matched with controls based on age, gender, race, and ethnicity. Our primary objective was to identify specific cell subsets and biomarkers associated with disease activity. Additionally, we aimed to assess the specific RA cell subsets and gene signatures in a diverse population, providing valuable insights into the multifaceted nature of RA pathogenesis.

## METHODS

### Study Design

RA Patients meeting the American College of Rheumatology (ACR) classification criteria (26) were recruited from the University of California San Francisco (UCSF) rheumatology clinic between 2016 and 2020 (27). Healthy controls were recruited through local advertising and through the database ResearchMatch(28). Controls were matched to RA patients by age, gender, race, and ethnicity. The study was conducted in accordance with the principles outlined in the Declaration of Helsinki and was granted ethical approval by the Human Research Protection Program and the Institutional Review Board of UCSF (IRB Project No. 15-17175). All participants provided written informed consent. Blood samples and participant level data were collected at time of enrollment. Clinical data including demographics, medication status, laboratory values, such as erythrocyte sedimentation rate, C-reactive protein (CRP), RF, ACPA antibodies, and clinical measurements of disease activity with the Disease Activity Score in 28 joints using CRP (DAS28-CRP). Patients were stratified in remission or low disease activity (DAS28-CRP <3.2) and moderate and high disease activity, (DAS28-CRP ≥ 3.2) according to the 2019 updated ACR recommendation on disease activity measures (29).

### Sample processing and 10X single cell RNA-sequencing

Peripheral Blood Mononuclear cells (PBMCs) were isolated from peripheral blood samples by UCSF Bay Area Center for AIDS Research Specimen Processing and Banking Subcore (previously AIDS specimen bank). Blood samples were collected in EDTA tubes, processed per manufacturer’s guidelines, and cryopreserved in liquid nitrogen. Single cell sequencing was performed using the 10X Chromium microfluidics system (10X Genomic). PBMCS from 18 RA patients and 18 healthy controls were thawed, counted, and pooled and profiled in three batches and 12 lanes, using 10x Genomics Chromium Single Cell 3’V3. Barcoded cDNA libraries were prepared using the single cell 3’mRNA kit. Cell Ranger v3 (3.1.0) was used to demultiplex cellular barcodes and map reads to the human (GRCh38-3.0.0) genome (30). Sample deconvolution and doublet identification was performed using demuxlet (31). RA and matched control samples were evenly split within each batch to limit technical and biological bias in our analysis. The first batch consisted of 14 individuals (seven RA and seven controls), the second batch included eight individuals (three RA and five controls, and the third batch included 16 individuals (eight RA and eight controls). One control was sequenced across three batches as a technical replicate to control for batch effect

### Preprocessing

Preprocessing was performed using Scanpy (1.9.1) (32) following previously published single cell workflows (33). Additional details on these methods can be found in our supplemental materials and published code. Genes found in less than three cells were filtered out, as well as cells containing fewer than 100 genes or more than 1000 genes. Cells with platelet or megakaryocyte gene markers (*PF4, GNG11, PPBP, SDPR*) were also removed. Additionally, cells containing greater than 20% mitochondrial genes, or less than 3% ribosomal genes were removed. Following filtering, the data were normalized to counts per million and log transformed. Technical variation from sequencing depth, mitochondrial percentage, ribosomal percentage were regressed out during scaling. Cell cycle scoring was performed using Scanpy using standard genes (34) and also regressed out. Batch correction was then performed using HarmonyPy (35), and samples were clustered in an unsupervised manner using leiden clustering with a resolution of 3.0. Each cluster was assigned as CD4+ T cells, CD8+ T cells, Monocytes, NK cells, or B cells using manual annotation with predefined marker genes according to the human protein atlas. Clusters of platelets, erythrocytes and suspected doublets were removed from further analysis. Sub-clustering was repeated for each cell type to allow for fine annotations of cell subsets, again based on referenced marker genes and leiden clustering. Highly expressed genes within each subset were identified using Wilcoxon rank testing implemented in Scanpy.

### Compositional analysis

Cell densities in each subset were calculated and plotted for RA samples versus controls using Scanpy embedding density functions. The proportion of each cell type within a sample relative to the total number of annotated cells for that sample was also calculated. Cells proportions were compared using Wilcoxon signed rank tests between RA and their matched controls. Additionally, Mann-Whitney U tests were used for cell proportions comparison among controls, RA patients with remission-low disease activity, and moderate-high disease activity. Correlation between cell-type proportion and DAS28-CRP was also performed by calculating Spearman rank-order coefficient. Two-sided p-value <0.05 was the threshold for statistical significance.

### Differential gene expression analysis

Differential gene expression analysis was performed between RA and matched controls using a pseudobulk approach using the bulk RNAseq tool DESeq2 (1.38.3) (36). Pseudobulk methods outperforms mixed models and limits pseudoreplication bias (37,38). For each cell subtype, read counts were summed across each sample to create a pseudobulk count matrix. DESeq2 was applied, using a Likelihood ratio test corrected on batch effect with an additional fit of a Gamma-Poisson Generalized Linear Model (GLM)(39). P-values were adjusted using Benjamini-Hochberg method and genes with a false discovery rate (FDR) ≤ 0.05 were selected. Additional filtering was applied with an absolute log2Fold change ≥ log2(1.6) and a base mean expression between 0.08 and 4.

### Over-representation analysis

Functional and over-representation analysis was performed on differentially expressed genes for each cell subtype using clusterprofiler (4.6.2) (40) and Gene Ontology (GO) database (41). We selected up and down regulated pathways related to biological processes, with a Gene Ratio ≥ 0.15, count ≥ 5 and (FDR) ≤ 0.01.

### Cell-cell communication analysis using CellChat

Cell-cell communication inference and visualization was performed using the CellChat R package (version 1.6.0) (42). CellChat uses the log-normalized expression matrix as input and predicts cell-cell communication based on ligand-receptor pairs in a curated database. For each pair of ligands and receptors, the communication probability is calculated based on the average expression of the ligand in one cell type and the average expression of the receptor in another cell type, considering the law of mass action. CellChat also considers other important signaling factors such as heteromeric complexes and cell type proportion in the estimation of the strength of interactions. We followed the standardized CellChat workflow, including the ‘projectData’-function, which allows for projecting the gene expression onto a validated protein-protein interaction network to impute the data. Cell-cell pairwise communication was visualized as the relative number of communications between groups of interest (RA v. control, low disease activity v. controls and high disease activity v. controls). Statistical significance (FDR adjusted p ≤ 0.05) of cell type sender and receivers was assessed by performing 50 permutations and comparing the results using a student’s t-test. Only pathways that were statistically significant (p ≤ 0.05) and with a relative contribution in the RankNet-function of either more than 0.65 or less than 0.35 were considered.

## RESULTS

### Experimental study design

We performed a single cell analysis on PBMC samples from 36 participants (18 RA and 18 controls matched on age, gender, ethnicity and race). Study design and population characteristics are described in **Fig. 1a, and Table 1**. The mean age was 53.75 ± 15.9 (mean ± standard deviation (SD)), the study was composed of 66.7% (N=24) of women, 11.1 % of Asian Americans, 83.3% of Caucasian, 5.6% of Latinx population. There was non-significant difference in population characteristics between RA patients and matched controls (**Table S1**). Clinical data regarding disease activity, antibody status, erosions and treatment were available for 16 RA patients. RA patients. Among the RA patients, 62.5% (N=10) presented a positive Rheumatoid Factor (RF), 87.5% (N=14) presented positive anti-citrullinated antibodies (ACPA). RA disease activity was evaluated using the Disease Activity Score on 28 joints using C-reactive protein (CRP) (DAS28-CRP) (29). The mean DAS28-CRP was 3.3 ± 1.0. Patients were stratified in remission - low disease activity (DAS28-CRP<3.2) (N=9) and moderate-high disease activity (DAS28-CRP ≥ 3.2) (N=7). Erosive disease was present in 62.5% of patients (N=10). The mean time since diagnosis was 4.13 ± 4.41 years. Seven patients were treated by conventional Disease-modifying antirheumatic drugs (DMARDs) (62.5%) and only one patient was treated by a biological DMARDs (TNF-inhibitor: etanercept).

**Figure 1.**
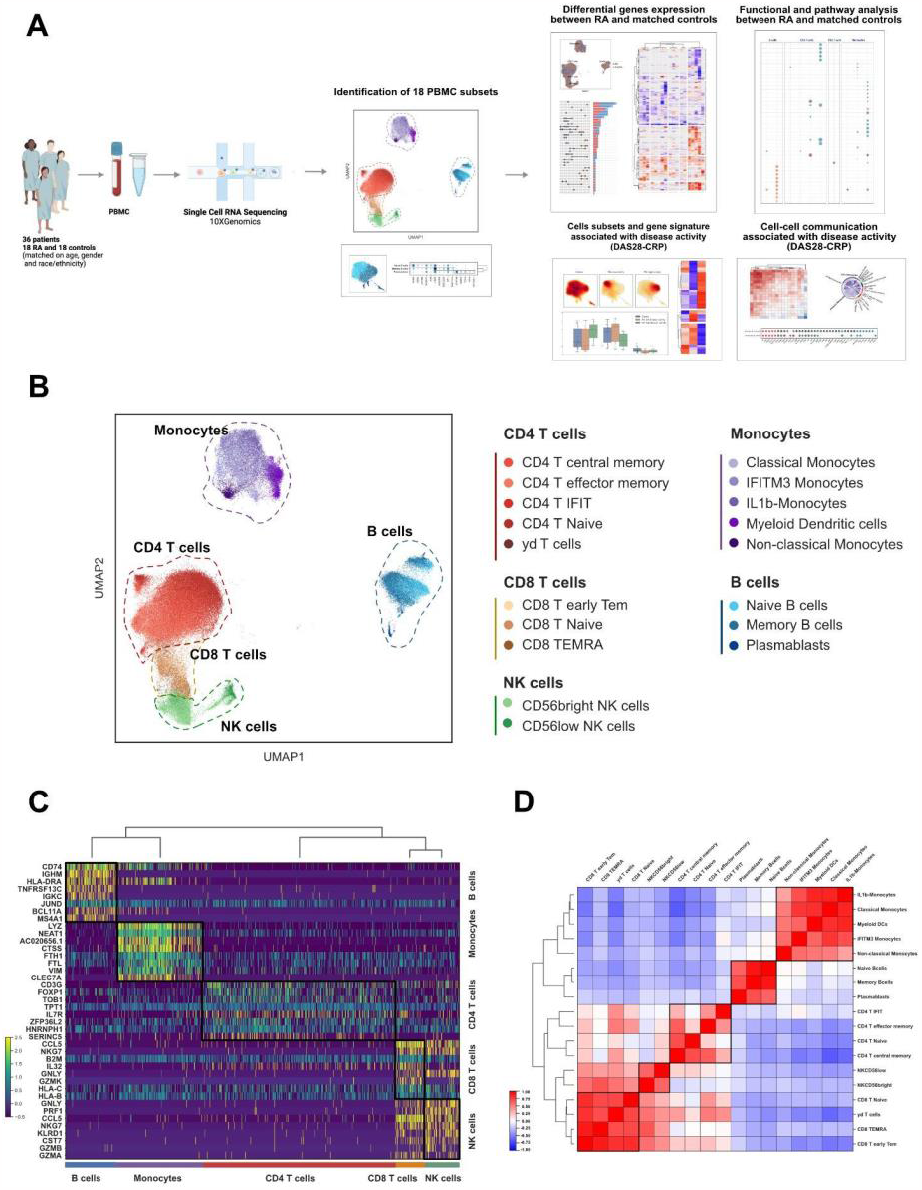
Identification of 18 PBMCs cell subsets including IFITM3+ Monocytes. A. Graphical Abstract : scRNA sequencing was performed on PBMCs from 36 individuals (18 RA and 18 matched controls on age, gender and race/ethnicity. Results identified cell subsets, differentially expressed genes using pseudobulk analysis, and cell-cell communication pathways enriched in RA versus controls, and in patients with high versus low disease activity .B. UMAP embeddings and subset annotations of single cell RNAseq dataset from patients with rheumatoid arthritis (n=18) and healthy controls (n=18) matched on age, sex, and ethnicity. C. Normalized expression of the top 40 ranked genes in different cell subsets (Wilcoxon rank test, FDR≤0.05). D. Correlation heatmap of genes expression across cells subsets (Spearman correlation). CD, Cluster differentiation; DCs, Dendritic cells; IFIT, Interferon Induced proteins with Tetratricopeptide repeats; IFITM, interferon-induced transmembrane; Tem, T effector memory, TEMRA, Terminally differentiated effector memory; RA, Rheumatoid Arthritis

**Table 1.** Clinical characteristics of RA patients and patients with remission-low and moderate-high disease activity. Disease activity information was available for 16 patients. p-value threshold for significance was set at <0.05. Student’s t-test was conducted to analyze continuous variables. For categorical variables, a Chi-square test was performed. ACPA, anti-citrullinated body mass index; bDMARDs: biological disease modifying antirheumatic drugs; cDMARDS, conventional disease modifying antirheumatic drugs; DAS: Disease activity score; RA, Rheumatoid Arthritis; RF, Rheumatoid Factor

|  |  | **Rheumatoid arthritis**<br>**(N = 18)** | **RA remission and low disease activity**<br>**(DAS28-CRP < 3.2)**<br>**(N = 9)** | **RA moderate and high disease activity**<br>**(DAS28-CRP ≥ 3.2)**<br>**(N = 7)** | **p-value** | **Missing values (%)** |
| --- | --- | --- | --- | --- | --- | --- |
| **Gender (%)** | Female | **12 ( 66.7 %)** | 7 ( 77.8 %) | 4 ( 57.1 %) | ns | 0 |
|  | Male | **6 ( 33.33)** | 2 ( 22.2 %) | 3 ( 42.9 %) |  |  |
| **Age (years) (mean (SD))** |  | **51.7 (15.3)** | 46.3 (15.4) | 59.2 (14.3) | ns | 0 |
| **Race (%)** | Asian | **2 ( 11.1 %)** | 1 ( 11.1 %) | 1 ( 14.3 %) | ns | 0 |
|  | Caucasian | **15 ( 83.3 %)** | 8 ( 88.9 %) | 6 ( 85.7 %) |  |  |
|  | Other | **1 ( 5.6 %)** | 0 ( 0.0 %) | 0 ( 0.0 %) |  |  |
| **Ethnicity (%)** | Hispanic | **1 ( 5.6 %)** | 0 ( 0.0 %) | 0 ( 0.0 %) | ns | 0 |
|  | Not Hispanic | **17 (94.4 %)** | 9 ( 100.0 %) | 7 ( 100.0 %) |  |  |
| **BMI kg/m2 (mean (SD))** |  | **26.7 (10.3)** | 24.41 (4.5 %) | 24.1 (5.4 %) |  | 5.6 |
| **DAS4 28 CRP (mean (SD))** |  | **3.3 (1.0)** | 2.6 (0.5) | 4.2 (0.6) | <0.001 | 11.1 |
| **RF (%)** |  | **10 ( 62.5)** | 4 ( 44.4 %) | 5 ( 71.4 %) | ns | 11.1 |
| **ACPA (%)** |  | **14 ( 87.50)** | 7 (77.8 %) | 6 (85.8 %) | ns | 11.1 |
| **Erosive status (%)** |  | **10 ( 62.5)** | 3 (33.3 %) | 5 (71.4 %) | ns | 11.1 |
| **cDMARDs (%)** |  | **7 (38.9 %)** | 5 ( 55.56) | 2 ( 33.33) | ns | 5.6 |
| **bDMARD (%)** |  | **1 (5.6 %)** | 0 (0 %) | 1 ( 14.3 %) | ns | 11.1 |
| **Disease duration (years)** |  | **4.13 (4.41)** | 3.61 (4.31) | 4.80 (5.22) | ns | 5.3 |

### Identification of 18 PBMC cell subtypes uncovering IFITM3+ monocytes

PBMCs collected on peripheral blood were pooled, profiled, and barcoded in three batches with 12 lanes using the 10x Genomics Chromium Single Cell technology. RA and matched controls were evenly split within each batch and lanes. Cell Ranger v3 was used for demultiplexing and read mapping to the human genome. Following 10X sequencing and preprocessing with Scanpy, our dataset consisted of 125,698 cells and 22,159 genes (**Fig. S1**). Leiden community detection was used to group cells into clusters, and annotation using established cell markers showed the presence of all major PBMC cell types (**Fig. S2**). All major cell types, annotated using established cell markers, were present in our dataset. Further clustering and annotation were used to identify cell subsets, including five CD4+ T cell subsets (CD4+ T central memory, CD4+ T effector memory, CD4+ IFIT+ T cells, CD4+ Naive T cells, gamma-delta (Vδ) T cells), three CD8+ T cell subsets (CD8+ Early T effector memory, CD8+ Naive T cells, Terminally differentiated effector memory (TEMRA)), two Natural Killer (NK) cell subsets (CD56bright NK cells, CD56 low NK cells), three B cell subsets (Naive B cells, Memory B cells and Plasmabasts), and five monocyte subsets (Classical Monocytes, IFITM3+, IL1-b, Myeloid dendritic cells, non-classical monocytes) (**Fig. 1b**). For the control sample that was replicated across batches, we found no statistically significant differences in cell proportions. Each of the cell subsets presented a distinct expression profile (**Fig. 1c, Fig. S2)**. CD4+, CD8+ T cells and NK cells subtypes showed higher similarity profiles, and the expression profiles of Vδ T cells exhibited a stronger correlation within the CD8+ T cells (**Fig. 1d**). Cell subsets and top genes of each cell-subset identified through Wilcoxon-rank sum analysis are included in **Fig. 2a-b**. We identified two cell subtypes (CD4+ IFIT+ cells and IFITM3+ monocytes) associated with genes related to Interferon (IFN)-pathway activation in RA. The pro-inflammatory CD4+ IFIT+ cell subtype presented significant expression of several genes associated with inflammatory response and immune regulation, including *IFIT2, PMAIP1, NFKBIZ, TNFAIP3* and *ZC3HAV1*. IFITM3+ monocytes had elevated levels of *IFITM3, ISG15, FTL, TYMP*, and *FTH1*. In addition, IFITM3 expression was specific to this monocyte subset (**Fig. S4**)

**Figure 2.**
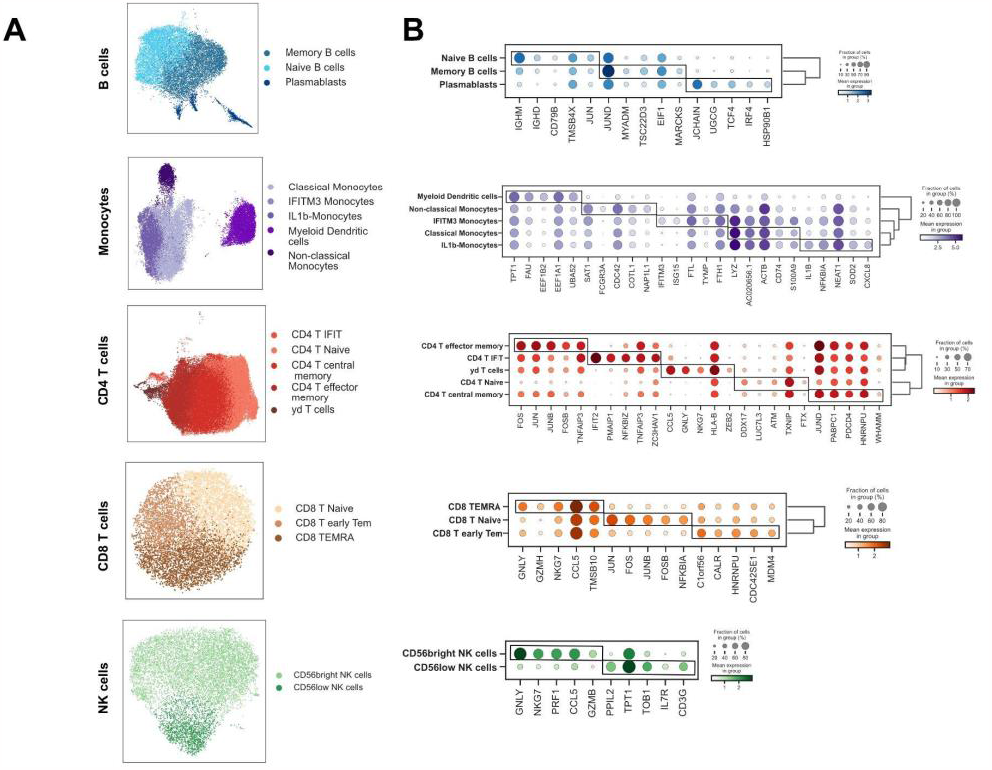
Cell subsets and top marker genes identified by Wilcoxon rank sum test. A. UMAP embedding for cell subsets in B cells, Monocytes CD4 T cells, CD8 T cells, NK cells. B. Dot plot of top ranking genes in each cell subset. CD, Cluster differentiation; DCs, Dendritic cells; IFIT, Interferon Induced proteins with Tetratricopeptide repeats; IFITM, interferon-induced transmembrane; Tem: T effector memory; TEMRA, Terminally differentiated effector memory; RA, Rheumatoid Arthritis

### Pseudobulk differential expression analysis revealed a specific down-regualtion of pro-inflammatory genes related to Vδ Tcells in RA patients in comparison to healthy controls

Non-classical monocytes cell proportions were significantly lower in RA patients compared to controls (Wilcoxon-signed rank analysis p=0.024) (**Fig. S3, Fig. S5**). By performing pseudobulk differential expression analysis on 18 cell subtypes, we identified a total of 168 genes that exhibited differential expression between individuals with RA and the control group (FDR ≤ 0.05, |log2FC| ≥ log2(1.6)). The majority of those genes were expressed in monocytes (n = 94) and CD8+ T cells (n = 39), 26 genes were expressed in CD4+ T cells, 6 in B cells, and 3 in NK cells (**Fig. 3a, 3b, Fig. 3c, table S2, table S3, Fig. S6)**. 121 genes were unique, and 47 genes were expressed across multiple cell types. Patients with RA had higher expression of genes associated with inflammation and cardiovascular risk in IL-1b and classical monocytes, including *IFITM2, TXNIP, EAF1, RIT1, EGR1, TLE3*, and *SLA*. In addition, they showed over-expression of cytotoxic genes *KLRD1, GZMH*, and *EBP* in CD8+ T cells. RA patients also displayed significant downregulation of proinflammatory genes such as *IFNG, IFIT2, TNF, GZMA, ISG15*, and *S100A4* exclusively in the Vδ T cells. Non-classical monocytes showed a specific transcriptomic profile of 19 differential expressed genes not shared with other cell-subsets **(Fig. 3b,c)**, including a downregulation of *ETNK1, TNFSF13B, DUSP7, IGSF6* and an upregulation of *CXCR4* in RA. Interestingly, the IFITM3+ subset also presented a cell-type specific downregulation of *HLA-DQB1, LRRK2, MS4A7*, and *G0S2* in RA.

**Figure 3.**
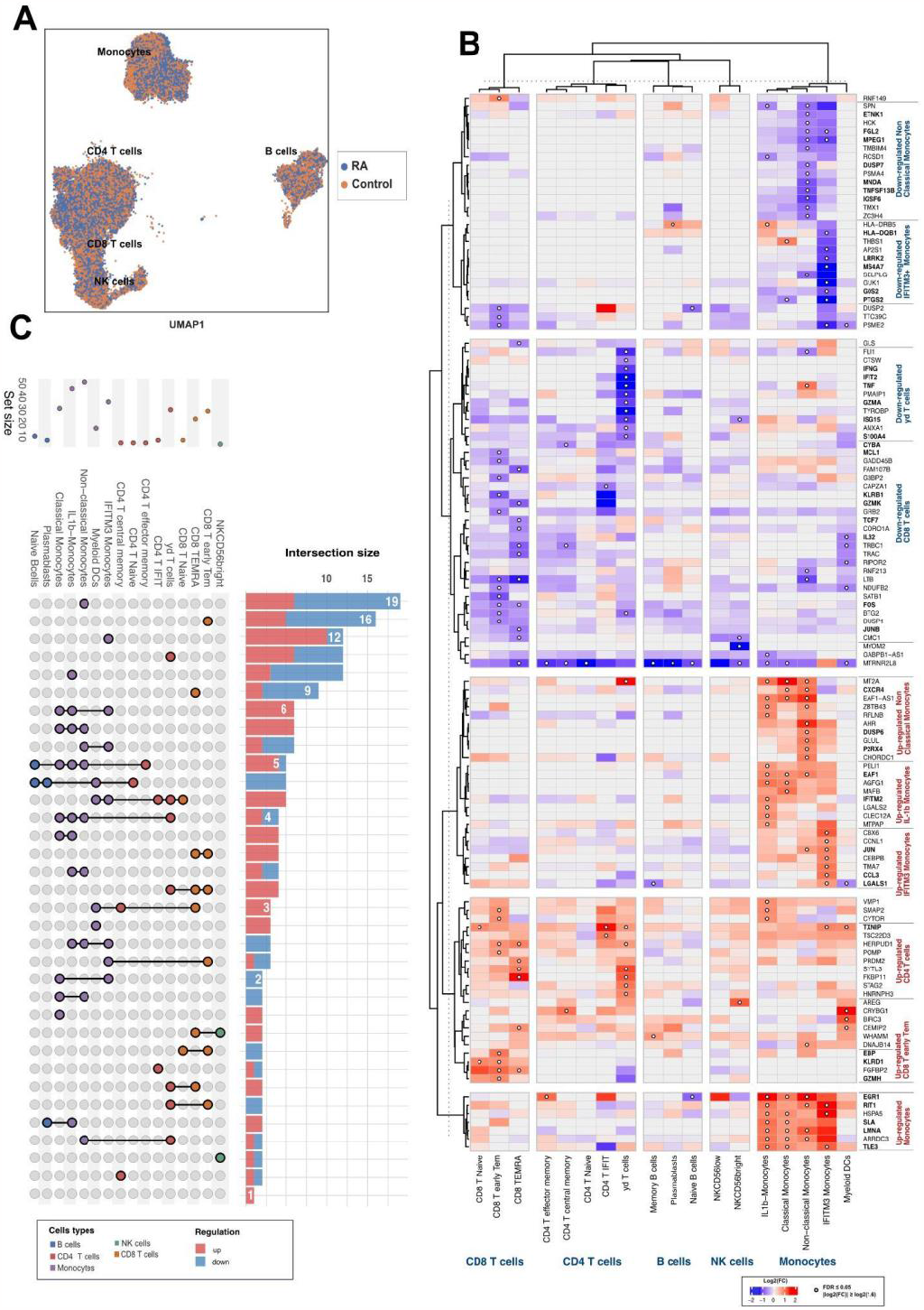
Pseudobulk analysis comparing RA patients and matched controls in each cell subset. A. Single cell UMAP projection of patients with rheumatoid arthritis and matched controls. B. Heatmap of differentially expressed genes between patients with RA and matched controls. C. Upset plot of upregulated and downregulated genes intersections across cell subsets.CD, Cluster differentiation; DCs, Dendritics cells; IFIT, Interferon Induced proteins with Tetratricopeptide repeats; IFITM, interferon-induced transmembrane; Tem, T effector memory; TEMRA, Terminally differentiated effector memory; RA, Rheumatoid Arthritis

### Over representation analysis found significant upregulation of B cell activation in RA patients

Functional analysis derived from pseudobulk differential expression analysis (FRD ≤ 0.05), identified 25 significantly up-regulated pathways across cell subsets in patients with RA compared to the control group. Among these pathways, 11 were up-regulated in B cells, 10 in Monocytes, and 4 in CD4+ T cells. Additionally, 21 pathways were significantly down-regulated, with 16 in Monocytes, 13 in CD4+ T cells, one in CD8+ T cells, and one in B cells (FDR ≤ 0.01, Gene ratio ≥ 15, counts ≥ 5) (**Fig. 4, Table S4**). We observed a noteworthy abundance of up-regulated pathways specifically in B cells of patients with RA as compared to the control group. These pathways primarily encompassed immune response, B cell activation, B cell receptor pathways, antigen-receptor mediated signaling pathways and immune response regulating cell surface receptor signaling pathways. Furthermore, within the Vδ T cell population, we observed a significant downregulation of pathways involved in positive regulation of myeloid and leukocyte differentiation as well as cytokine production regulation, in RA patients. Finally, in non-classical monocytes, we observed an up-regulation of cytokine-mediated signaling pathway and a downregulation of pathways involved in T cell activation, lymphocytes regulation and mononuclear cell proliferation, and leukocyte cell-cell adhesion in RA patients.

**Figure 4.**
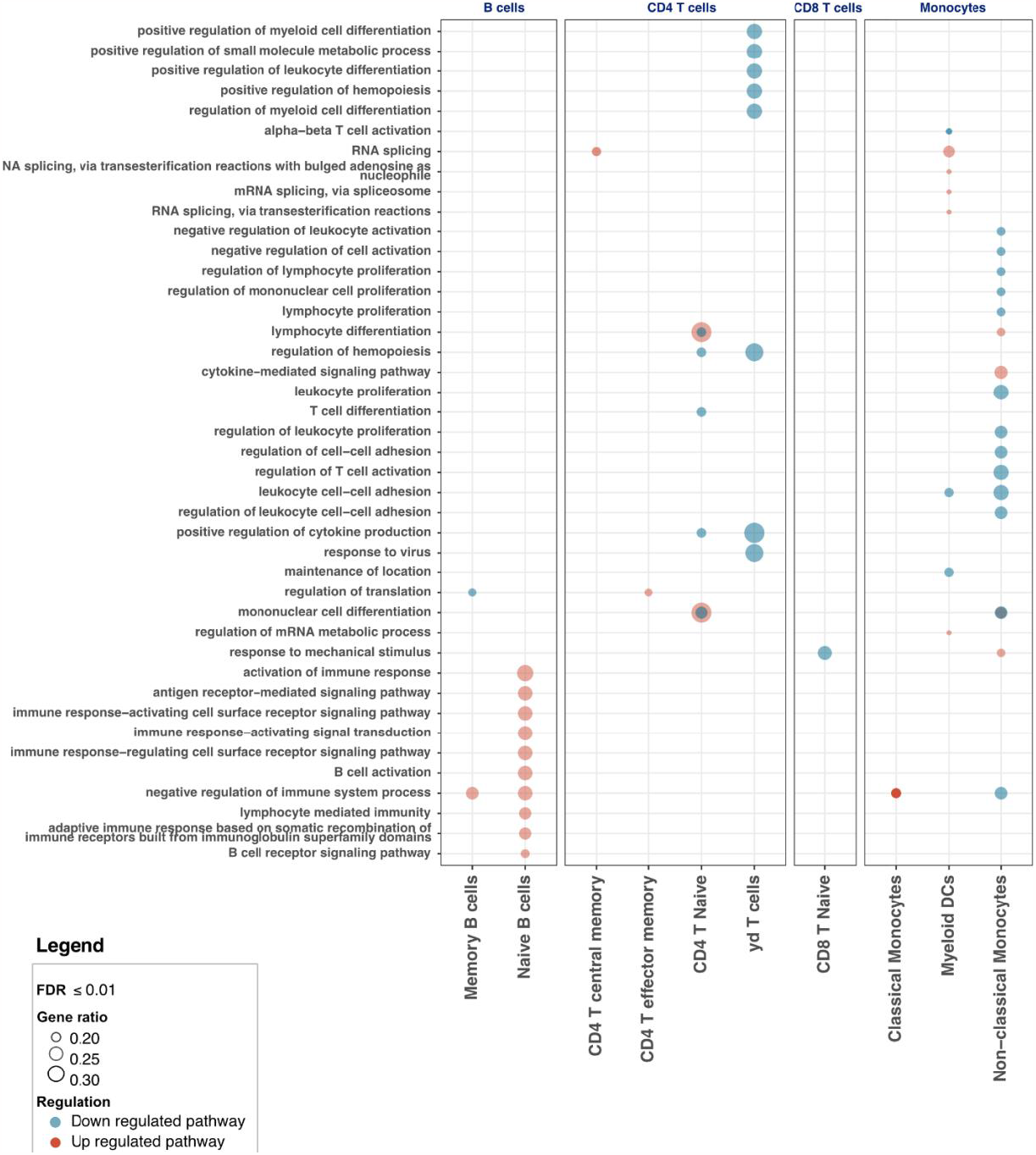
Functional analysis comparing RA and matched controls. Significant pathways from over-representation analysis, across 18 cells subtype (Gene ratio > 0.15, FDR ≤ 0.05, 0.08 < base mean < 4). CD, Cluster differentiation; DCs, Dendritic cells; IFIT, Interferon Induced proteins with Tetratricopeptide repeats; IFITM, interferon-induced transmembrane; Tem, T effector memory; TEMRA, Terminally differentiated effector memory; RA, Rheumatoid Arthritis

### CD4+ central memory cell and Non-classical monocytes proportions were associated with disease activity in RA patients

We performed a stratified analysis based on disease activity comparing individuals with active vs. inactive disease. Information regarding DAS4-28-CRP was available in 16 out of 18 RA patients included in the study. RA patients were divided into two groups: one consisting of individuals in remission or with low disease activity, characterized by a DAS28-CRP <3.2 (N=9), and the other group comprising patients with moderate and high disease activity, indicated by DAS28-CRP ≥3.2 (N=7). There was no significant statistical difference between the two groups in terms of age, gender or sex, race, ethnicity, BMI, proportions of ACPA and RF, erosive disease, treatment strategy, and disease duration **(Table 1**). We compared differences in cell density and cell subset proportions between those two groups and the control group (**Fig. 5a-c**). Additionally, we conducted a non-parametric Spearman correlation analysis to evaluate the association between cell proportions and DAS28-CRP as a continuous score (**Fig. S7**). Although there was no statistical significance in B cell proportion across groups, we observed a clear density shift between patients with remission low disease activity vs moderate and high activity from Naive to activated Memory B cells (**Fig. 5b**). RA patients with moderate to high disease activity showed a significantly increased proportion of CD4+ central memory cells (p=0.034) (**Fig. 5c**). Conversely, Non-classical monocytes were significantly lower in patients in the remission-low disease activity group compared to both control and the group with moderate-high disease activity (p=0.022).

**Figure 5.**
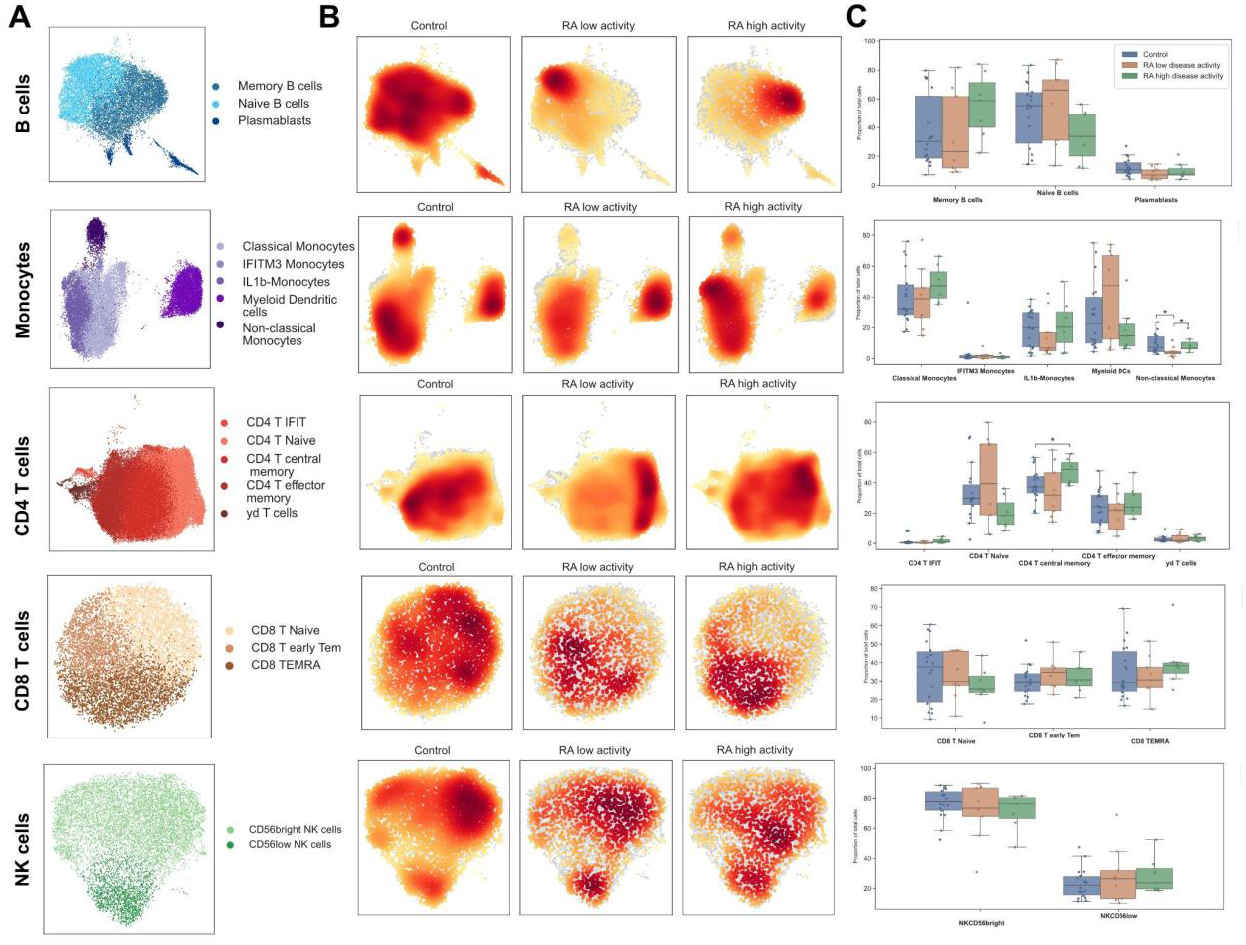
Cell proportion and cell density between patients with low and high disease activity and matched controls A. UMAP representation of cell subsets. B. Compositional and density plots comparing control patients with low disease and high activity. C. Cell proportion analysis between controls and RA patients with low disease and high activity. (Mann-Whitney - Wilcoxon p≤ 0.05). CD, Cluster differentiation; DCs, Dendritic cells; IFIT, Interferon Induced proteins with Tetratricopeptide repeats; IFITM, interferon-induced transmembrane; Tem, T effector memory; TEMRA, Terminally differentiated effector memory; RA, Rheumatoid Arthritis

### Identification of a gene signature specific to moderate and high disease activity in RA

Using a gene list consisting of 121 unique genes that exhibited differential expression between individuals with RA and matched controls in our pseudobulk analysis, we conducted a subanalysis focusing on patients with different disease activity levels: those in remission or with low disease activity, and those with moderate to high disease activity. Among the 121 genes, 75 were significantly associated with low disease activity, and 89 associated with high disease activity (FDR ≤ 0.05, |log2(FC)| ≥ log2(1.6), 0.08 ≤ base, mean < 4). Interestingly, 52 genes were significantly upregulated in patients within the moderate to high disease activity group. These genes included *G0S2, THBS1, DUSP7, IFIT2, IGSF6, MAFB, RIT1, TNF, JUN, CXCR4*, and *TLE3* (**Fig. 6, Fig. S8**). Furthermore, we observed a separate set of 37 genes that exhibited a significant downregulation in patients with moderate to high disease activity compared to the control group. These downregulated genes included *TRBC1, KLRB1, IL32, HLA-DQB1, HLA-DRB5, TNFSF13B, CCL3, LRRK2*, and *TMA*7. Additionally, we identified a set of 12 genes that were significantly overexpressed only in patients with remission or low disease activity, which included *TXNIP, LGALS2*, and *AREG*.

**Figure 6.**
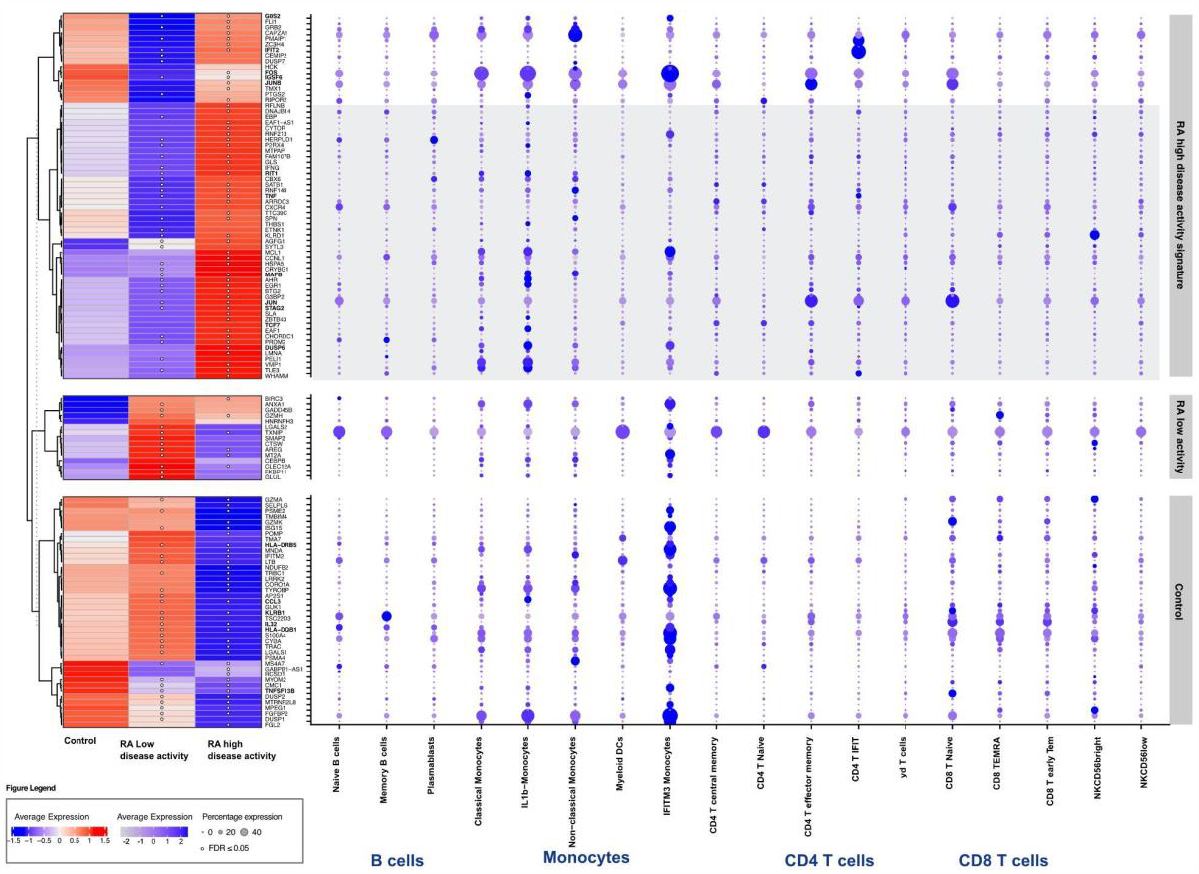
Gene signatures associated with disease activity across cell subsets. Gene expression heatmap comparing controls and RA low and high disease activity adjacent to a dot plot showing average expression across cell subtypes. CD, Cluster differentiation; DCs, Dendritics cells; IFIT, Interferon Induced proteins with Tetratricopeptide repeats; IFITM, interferon-induced transmembrane; Tem, T effector memory; TEMRA, Terminally differentiated effector memory; RA, Rheumatoid Arthritis

### CD4+ T cells and B cell subsets were associated with the highest levels of cell-cell communication in RA patients

To gain a comprehensive understanding of immune cell communications, we conducted a cell-cell communication inference analysis using CellChat, which uses a repeated permutation to identify significant cell-cell interactions. We found a statistically significant increase in cell-cell communication in RA patients as compared to healthy controls in 35 pairs of cell types. The largest increase in communication was found in CD4+ naive T and CD4+ T central memory cells along with Naive and Memory B cells both as senders and receivers. In addition, there was an increase between Classical, Non-classical and IL-1β Monocytes as senders and Naive CD4+ T cells as receivers. One cell-cell pair showed no difference, and 255 cell-pairs presented less communication in RA (**Fig 7a**) When stratifying patients based on disease activity, we found similar patterns. In patients with remission-low disease activity, 28 pairs were significantly increased, and 286 pairs were decreased in communication as compared to controls **(Fig S9**). In patients with moderate and high disease activity, 37 pairs had statistically significant increases in communication and 259 pairs presented with significant decreases in communication, while one pair had a similar amount of communication as compared to controls. For the high activity states, decrease as both sender and receiver were observed in particular for CD8+ T naive and early TEM and CD4+ T effector memory as well as CD56 bright NK cells and IFITM3 monocytes, whereas the remission-low disease activity state presented a decrease of both sending and receiving communication of IFIT+ CD4+ T cells, myeloid dendritic cells as well as IFITM3 monocytes. On the other hand, Myeloid dendritic cells appeared to be more involved in communication in high disease activity as both a sender and receiver, while NK cells and CD4+ naive T cells were more involved in sending and receiving communication in low disease activity.

**Figure 7.**
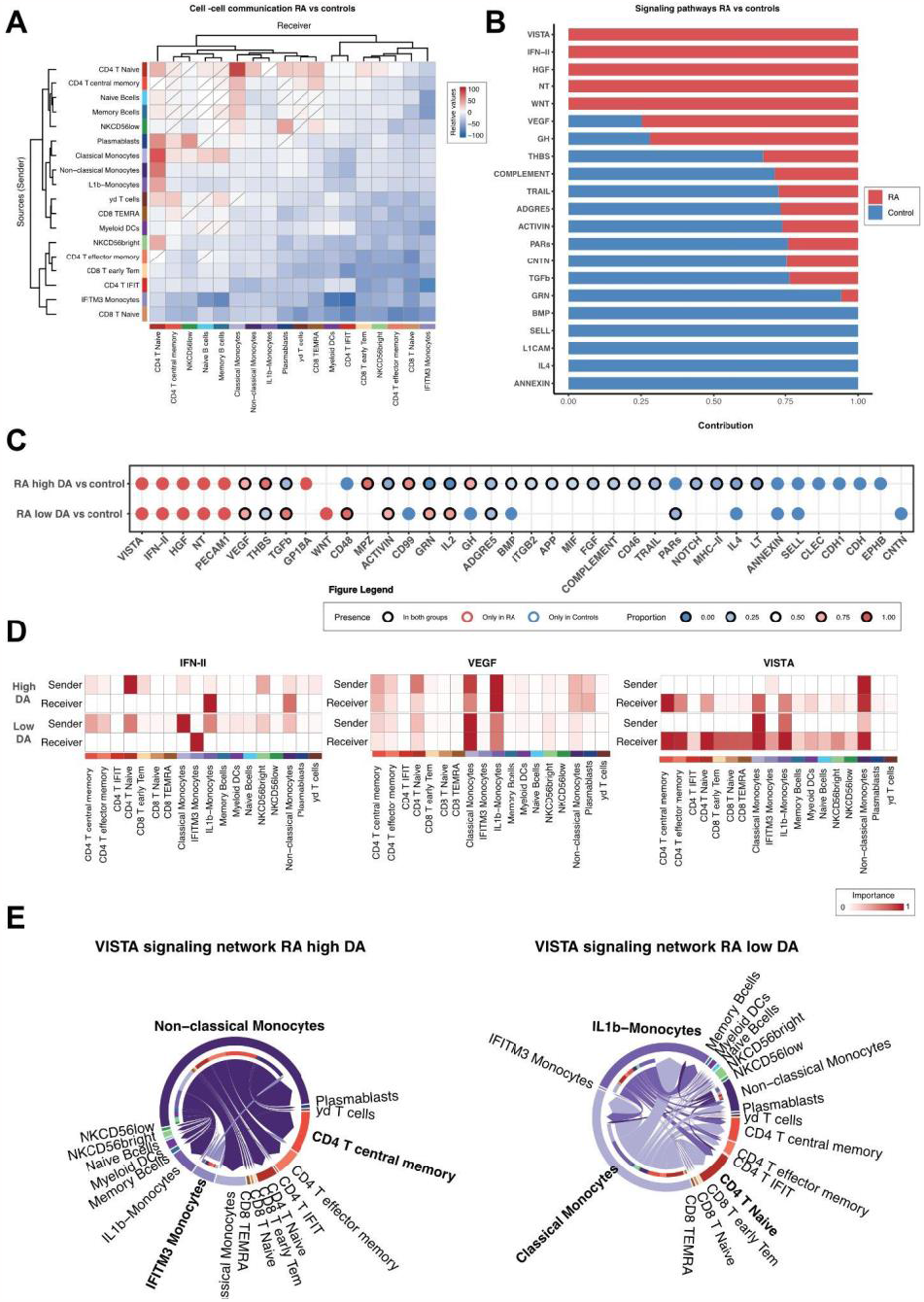
Cell-cell communication patterns between patients with low and high disease activity and matched controls. A. Heatmap representing the relative number of interactions between RA and matched controls. B. Barplot of statistically significant communication pathways based on strength of interactions between RA patients and controls. C. Dot plot of the relative contributions of communication pathways based on strength of interactions between high and low disease activity compared to controls. D. Heatmaps of the relative importance of cells as senders and receivers for the IFN-II, VEGF and NT signaling pathways in high and low disease activity. E. Circle plots representing the relative importance of cells as senders and receivers for the VISTA signaling pathway network in high and low disease activity and overall. CD, Cluster differentiation; DA: disease activity; DCs, Dendritics cells; IFIT, Interferon Induced proteins with Tetratricopeptide repeats; IFITM, interferon-induced transmembrane; Tem, T effector memory; TEMRA, Terminally differentiated effector memory; RA, Rheumatoid Arthritis

### Cell-cell communication revealed an upregulation of VISTA and Interferon II pathways in RA patients with moderate and high disease activity

CellChat utilizes a curated database of ligand-receptor pairs, grouped into communication pathways which may contain multiple ligand-receptor pairs. The change in communication pathways between disease states is then calculated as the relative contribution per disease state to the total communication amount for a specific communication pathway. We used a threshold for significant contribution at less than 35% or more than 65% of the total communication and a p-value < 0.05. We found several distinct communication pathways to be up- or downregulated in RA, with 7 pathways showing more communication and 14 showing less communication as compared to controls **(Fig 7b, Fig S10)**. Upregulated pathways included NT, HGF, VISTA, IFN-II and WNT. When comparing moderate-high disease activity to low disease activity, we found the highest number of changed pathways to be in moderate-high (36 vs 22) (**Fig 7c, Fig S11a-c**). Many pathways from the initial comparison remained significantly and solely expressed in RA, including NT, HGF, VISTA and IFN-II, although PECAM-1 also became significant. For each pathway, specific ligand-receptor pairs were considered main contributors (**Fig S12)**.

For most pathways, the direction of dysregulation was similar in both high and low disease activity (n = 12). However, the THBS (thrombospondin), CD99 and GH pathways were upregulated in high disease activity but downregulated in low disease activity. On the other hand, there was more communication in TGFb, activin, GRN and IL-2 pathways in low disease activity as compared to high disease activity. Although the direction of dysregulation was alike for high and low disease activity, the cells contributing to communication pathways differed. For IFN-II, naïve CD4+ T cells and CD56 bright NK cells sent signals to primarily IL1b and nonclassical monocytes in high disease activity (**Fig 7d**). In low disease activity, most signaling came from classical and IL-1b monocytes and was received by IFITM3 monocytes (**Fig 7d**). In other pathways, such as VEGF and NT, the involved cells remained unchanged across disease states; however, small changes in importance were observed (**Fig 7c-d, Fig S13**). For IL2, most communication was between CD4+ cells, although myeloid dendritic cells also played a large role in high disease activity (**Fig S13**) Another upregulated pathway across disease states was the VISTA pathway, for which we found discrepancies in the involved cells, as non-classical and IFITM3 monocytes were the primary senders in high disease activity as opposed to low disease activity where non-classical monocytes were outshadowed by classical and IL1B-monocytes (**Fig 7e**). For both disease states; however, the main receivers were within the CD4+ and myeloid subsets, although CD4+ Naive T cells were more prominent receivers in low disease activity, and CD4+ Memory T cells were more prominent in high disease activity.

## DISCUSSION

Here we describe a dataset of scRNA-Seq of PBMCs from a diverse population of 36 patients with 18 RA and 18 matched controls on age, gender, race and ethnicity.

In our study we identified a monocyte subset specifically expressing IFITM3+. IFITM3, also known as IFN induced trans-membrane 3, is associated with type 1 IFN response and viral restriction (43). Associations between IFITM3 haplotypes and rheumatoid arthritis have been reported in the literature, particularly in the context of a Korean population (44). IFITM3 polymorphism has also been associated with other auto-immune diseases such as ulcerative colitis (45). By identifying a IFITM3+ monocyte subset in the blood our results reinforce the potential role of IFITM3 in RA pathophysiology and especially in monocytes. Our results also reinforce previous findings by Zhang et al. who identified through the integration of mass cytometry and bulk RNA and upregulation of *IFITM3* genes associated with monocytes subsets in synovial tissue(17). Interestingly several genes associated with RA genetic predisposition were also differentially expressed in the IFITM3+ monocytes subsets such as *HLA-DQB1, LRRK2, TLE3. LRRK2* mutations have been associated with several auto-immune diseases and especially rheumatoid arthritis predisposition (46,47). In addition, a meta-analysis demonstrated that *HLA-DQB1* polymorphisms were associated with RA with a protective role of DQB1*02 and DQB1*06 and conversely a susceptibility role of DQB1*04 (48). In addition, a variant in *TLE3* has been associated with ACPA-positive rheumatoid arthritis in several European populations(49).

In our pseudobulk differential expression analysis, we observed a specific down regulation of pro-inflammatory genes in the Vδ T cells subsets including, *IFNG, IFIT2, TNF, GZMA, ISG15, S100A4*. Interestingly Mo et al. described that peripheral Vδ2T cells were significantly lower in patients with RA and were negatively correlated with disease activity. In addition, they described that *V*δ2 T cells from RA accumulated in the synovium and produced high levels of pro-inflammatory cytokines including IFN-γ and IL-17 and also showed elevated chemotaxis potential (50). Our results could reinforce the potential chemotaxis role of *V*δ2 T cells in the synovium. However, it is essential to acknowledge that this remains a hypothesis as we lacked matched tissue data in our study to confirm this hypothesis conclusively. Similarly, we observed a lower proportion of non-classical monocytes in PBMCs from patients with remission-low disease activity groups compared to both controls and moderate-high disease activity. Guła et al have reported that the absolute number of circulating non-classical monocytes negatively correlates with DAS28 and swollen joint count in peripheral spondyloarthritis patients (51). Non-classical monocytes have also been associated as key mediators of tissue destruction in osteoclasts in murine models of rheumatoid arthritis(52). In addition we found a specific downregulation in the non-classical monocytes subsets of several genes such as *IGFS6, ETNK1, DUSP7*, and *TNFSF13B. IGFS6* expression has been significantly associated with RA fibroblast cells in humans(53) *Etnk1* has been associated as a candidate gene in collagen induced arthritis(54). *DUSP7* is involved in MAPK signaling and low levels of mRNA of *DUSP7* have been associated with RF ACPA positive RA patients(55). *TNFSF13B* variants have been associated in several studies with RA and also with other auto-immune diseases such as systemic erythematous lupus(56,57)

We also identified a gene signature of 89 genes specific to disease activity including an upregulation of proinflammatory genes such as *TNF, JUN, EGR1, IFIT2, IGSF6, TMX1*, and *MAFB*, but also potential genes associated as therapeutic targets or in treatment response such as *G0S2, PTGS2*, and *THBS1. Mafb* has been associated with monocytes and macrophage differentiation but also involved in the activation of myeloid cells associated with joint destruction such as RANK+ TLR2-cells in murine models of RA (58,59). *G0S2* has been associated with anti-TNF response prediction in a meta-analysis of 11 studies(60). Thrombospondin-1 expression has been associated with NR4A2 activity and is modulated by TNF inhibitors (61). We also observed several genes downregulated in patients with high disease activity such as *HLA-DQB1, HLA-DRB5*, and *TNFSF13B*. Similarly, Klimenta et al. also found a protective role of HLA-DRB5 in RA (62). In an independent single cell study, Wu et al. showed that HLA-DRB5+ expression was lower in the synovial tissues of ACPA-RA patients(18).

Cell-cell communication allowed us to confirm several well-known signaling pathways from the RA literature, including type II IFN (IFN-γ), TGFb and VEGF(63–65). Additionally, our findings revealed an upregulation of the IL-2 signaling pathway in patients with remission and low disease activity, whereas a downregulation was observed in patients with moderate and high disease activity compared to controls. Those paradoxical findings align with previous research by Tebib et al., who also reported lower IL-2 levels in patients with active disease compared to those in a non-active state, although the difference did not reach statistical significance (66). IL-2 has been correlated with disease activity and severity in several studies (66,67). The IL-2 pathway plays a pivotal role in Regulatory T cell response and holds significance in rheumatic diseases (68). Furthermore, the concept of utilizing low-dose IL-2 as a potential therapeutic target in RA has been proposed (69–71). Consequently, gaining a deeper understanding of the role of IL-2 is imperative. However, it should be acknowledged that our study lacked sufficient power to identify specific regulatory T cell subsets and did not identify IL-2 in our differential expression gene list. Thus, further research is warranted to refine and enhance this hypothesis.

We also found an up-regulation of the VISTA signaling pathway in RA patients. VISTA is a negative checkpoint regulator, playing a key role in suppressing T cell-mediated immune responses, and its disruption has been linked to pro-inflammatory phenotypes and a susceptibility to autoimmune diseases (72). In a recent groundbreaking study, ElTanbouly et al. investigated the role of VISTA expression in T cells and found that VISTA expressed on naive T cells was playing a critical role in quiescence and peripheral tolerance (73). In our signaling network analysis of VISTA, CD4+ T cells subsets were the main receivers. CD4+ naive T cells were the primary receiver population in the remission-low disease activity group, while CD4+ T central memory was the predominant group in moderate-high disease activity. Interestingly, the monocytes cells population were associated as the main sender in the VISTA signaling network, in particular IL-1β Monocytes and classical monocytes in remission-low disease activity and Non-classical and IFITM3 monocytes in moderate and high disease activity groups. A study conducted on human PBMCs revealed that VISTA expression predominates in monocytes, and was associated with CD11b cells in murine models(74). Furthermore, previous results from Ceeraz et al have found a significant reduction of arthritis score in VISTA deficient mice and mice treated with monoclonal anti-VISTA antibodies in a collagen antibody-induced arthritis model independent of T- and B-cells, supporting our finding of the importance of innate, myeloid-derived VISTA in arthritis(72). Targeting VISTA shows promising potential as an innovative immunoregulatory therapy (75). Our findings further support the involvement of VISTA in RA and its potential impact on the communication between monocytes and CD4+ T cells. However, additional research is necessary to gain a comprehensive understanding of VISTA’s role in RA.

Several limitations should be acknowledged in our study. First, it is challenging to use scRNA-seq of unsorted PBMCs to study very small cell subsets, such as regulatory T cells, which may have led to their role in disease activity being underestimated and not extensively explored. Additionally, the relatively small sample size of our study may have limited the statistical power to detect subtle differences. This limitation also restricted our ability to thoroughly explore potential differences associated with gender, race, and ethnicity, emphasizing the need for more inclusive representation in future investigations to ensure a comprehensive understanding of RA across diverse populations.

Another limitation is the absence of matched disease synovial tissue, which could have provided more comprehensive insights into how patterns in PBMCs relate to cellular and molecular mechanisms involved in RA synovium.

Nonetheless, our study provides valuable insights into the cellular and molecular mechanisms associated with disease activity in RA. We carefully considered matched controls on age, gender, race, and ethnicity, and our work has identified key cell subsets and genes that may be associated with disease activity. These findings have the potential to serve as new biomarkers and therapeutic targets. We are optimistic that our research will contribute to advancing our understanding of RA pathogenesis and lead to the development of more effective treatments for this complex autoimmune disease.

## Acknowledgements

We would like to thank the patients who contributed to the study and the administrative and medical team of UCSF rheumatology clinics. We would like to thank the PREMIER center as well as members of the Sirota Lab for useful discussion. This work is in part supported by Pfizer and NIAMS P30 AR070155.

## Author’s contribution

AG and LAC recruited the patients for the study, MN, EN performed the single cell experiments, D.R. and W.T. performed the alignment and the samples deconvolution. M.B. and B.Y.M. performed the quality controls, the pre-processing and the cell proportion analyses. M.B., B.Y.M. and C.W. contributed to the cell-type annotations. M.B. performed the pseudobulk analysis and the analysis associated with disease activity. M.B. and C.W. performed the functional analysis. C.W. performed the cell-cell communication analysis. E.F. and U.K. peer-reviewed and provide essential feedback on the code and the analysis strategy. M.Y. and M.N. provided complementary information on the study metadata. M.B., B.Y.M., C.W., and M.Y. wrote the manuscript and M.S. revised the manuscript. M.S, L.A.C and M.N conceptualized the study and M.S was involved in the funding acquisition. M.S was involved in the project administration. Y.S, G.F, J.N, J.S, D.K. E.M.F A.J.B, A.G, J.Y, L.A.C M.N provided critical feedback and edits to the manuscript.

## Funding

This work was supported by the Rheumatology Research Foundation Targeted Research Pilot Grant p30 from the National Institutes of Health (NIH), National Institute of Arthritis and Musculoskeletal and Skin Diseases (NIAMS) (P30-AR070155) and additional funding from Pfizer.

## Key messages

1. *What is already known on this topic* - Previous transcriptomic studies have identified genes associated with rheumatoid arthritis (RA) in synovial tissue and peripheral blood, but the molecular and cellular signatures associated with heterogenous RA disease activity remain poorly understood.
2. *What this study adds* – Here, we characterize the single cell transcriptomic profiles of peripheral blood mononuclear cells in patients with RA and healthy controls, and provide insights into differentially expressed genes and cells associated with disease and disease activity
3. *How this study might affect research, practice or policy* – The identification of cellular and molecular signatures in patients with RA, particularly markers associated with disease activity, can provide valuable insights into novel targets for drug discovery or effective management of RA disease.

## Data and materials availability

Upon manuscript acceptance, the data and code will be made publicly accessible on GO, dbGap, and GitHub

## Supplementary Materials

Fig. S1 to S13

Table S1 to S2

## Notes

### Competing Interest Statement

MB research is funded by a grant from the French Society of Rheumatology, the Osteoarthritis Foundation grant and a doctoral fellowship from Sorbonne University. BYM is a paid consultant for SandboxAQ. CW research is funded by grants from the Lundbeck Foundation, Fabrikant Aage Lichtingers Foundation, Director Ib Henriksens stipend, Knud Hoejggaards Foundation, Jorcks Foundation and the Denmark-America Foundation. CW is a minor share holder in Ambu A/S, Bavarian Nordic A/S, Genmab A/S, H. Lundbeck A/S A and B and several other non-health related companies and mutual funds. U.K. is a paid consultant for Vevo Therapeutics. AJB is a co-founder and consultant to Personalis and NuMedii, consultant to Samsung, Mango Tree Corporation, and in the recent past, 10x Genomics, Helix, Pathway Genomics, and Verinata (Illumina), has served on paid advisory panels or boards for Geisinger Health, Regenstrief Institute, Gerson Lehman Group, AlphaSights, Covance, Novartis, Genentech, and Merck, and Roche, is a shareholder in Personalis and NuMedii, is a minor shareholder in Apple, Facebook, Alphabet (Google), Microsoft, Amazon, Snap, 10x Genomics, Illumina, CVS, Nuna Health, Assay Depot, Vet24seven, Regeneron, Sanofi, Royalty Pharma, AstraZeneca, Moderna, Biogen, Paraxel, and Sutro, and several other non-health related companies and mutual funds, and has received honoraria and travel reimbursement for invited talks from Johnson and Johnson, Roche, Genentech, Pfizer, Merck, Lilly, Takeda, Varian, Mars, Siemens, Optum, Abbott, Celgene, AstraZeneca, AbbVie, Westat, and many academic institutions, medical or disease specific foundations and associations, and health systems. Atul Butte receives royalty payments through Stanford University, for several patents and other disclosures licensed to NuMedii and Personalis. Atul Buttes research has been funded by NIH, Peraton (as the prime on an NIH contract), Genentech, Johnson and Johnson, FDA, Robert Wood Johnson Foundation, Leon Lowenstein Foundation, Intervalien Foundation, Priscilla Chan and Mark Zuckerberg, the Barbara and Gerson Bakar Foundation, and in the recent past, the March of Dimes, Juvenile Diabetes Research Foundation, California Governors Office of Planning and Research, California Institute for Regenerative Medicine, LOreal, and Progenity. None of these entities had any role in the design, execution, evaluation, or writing of this manuscript.

