## Supplemental figures and tables for "Deciphering Cell-types and Gene Signatures Associated with Disease Activity in Rheumatoid Arthritis using Single Cell RNA-sequencing"

**Figure S1 | Quality control and batch correction with HarmonyPy A. UMAP projection according to sample type (control or RA) batch and lane before batch correction with Harmony. B. UMAP projection according to sample type, batch and lane after batch correction with Harmony. C. UMAP projection of lanes after batch correction with Harmony.** RA, Rheumatoid Arthritis; UMAP, Uniform Manifold Approximation and Projection

**Figure S2 | Expression heatmap representing the mean expression of B cells, CD4 T cells, CD8 T cells, NK cells and Monocytes marker genes for cell subsets.** DCs, Dendritics cells; Tem, T effector memory; TEMRA, Terminally differentiated effector memory

**Figure S3 | Cell proportions of B cells, CD4 T cells, CD8 T cells, NK cells and Monocytes across samples.** CD, Cluster Differentiation; NK, Natural Killer; RA, Rheumatoid Arthritis

**Figure S4 | A. UMAP representation of IFITM3 gene expression across all PBMCs cell types. B. UMAP visualization of IFITM3 gene expression across monocyte subsets.** CD: Cluster Differentiation; DCs: Dendritic Cells; IFIT: Interferon Induced proteins with Tetratricopeptide repeats; IFITM: Interferon-induced Transmembrane proteins; Tem: T Effector Memory; TEMRA: Terminally Differentiated Effector Memory. RA: Rheumatoid Arthritis

**Figure S5| A. Cell subsets for B cells, Monocytes, CD4 T cells, CD8 T cells, NK cells. B, Compositional analysis and density plots comparing patients with Rheumatoid Arthritis and matched controls. C. Cell proportion analysis between patients with Rheumatoid Arthritis and Controls (Wilcoxon signed rank test $p \leq 0.05$).** CD, Cluster differentiation; DCs, Dendritic cells; IFIT, Interferon Induced proteins with Tetratricopeptide repeats; IFITM, interferon-induced transmembrane; Tem, T effector memory; TEMRA, Terminally differentiated effector memory; RA, Rheumatoid Arthritis

**Figure S6 | Volcano Plot representing the differential gene expression analysis between patients with Rheumatoid Arthritis and matched controls in each cell subset ($FDR \leq 0.05$, $\log2(FC) \geq \log2(1.6)$, $0.08 \leq$ mean expression$< 4$ ).** CD, Cluster differentiation; DCs, Dendritic cells; FC, Fold change; FDR, False discovery rate; IFIT, Interferon Induced proteins with Tetratricopeptide repeats; IFITM, interferon-induced transmembrane; Tem, T effector memory; TEMRA, Terminally differentiated effector memory; RA, Rheumatoid Arthritis

**Figure S7 | A. Spearman correlation of cell subset proportion with the DAS28-CRP for each subset. B Correlation heatmap of cell subset and DAS-28-CRP.** CD, Cluster differentiation; DC, Dendritic cells, IFIT, Interferon Induced proteins with Tetratricopeptide repeats; IFITM, interferon-induced transmembrane; Tem, T effector memory; TEMRA, Terminally differentiated effector memory.

**Figure S8 | Venn Diagram of the genes differentially expressed in RA low and high disease activity compared to controls. ($FDR \leq 0.05$, $\log2(FC) \geq \log2(1.6)$, $0.08 \leq$ mean expression $< 4$ ).** RA, Rheumatoid Arthritis

**Figure S9 | Comparisons of cell-cell communication patterns between patients with low and high disease activity and controls. A. Heatmap representing the relative number of interactions between cell types of RA patients with high disease activity compared to controls. B. Heatmap representing the relative number of interactions between cell types of RA patients with low disease activity compared to controls.** DA, Disease activity

**Figure S10 | A. Bar plots of all communication pathways based on interaction strength (red are significantly more present in RA, blue are significantly more present in controls). B. Bar plots of all communication pathways based on number.** DA, Disease Activity; RA, Rheumatoid Arthritis

**Figure S11 | A. Dot-plot of the relative contribution of communication pathways based on number of interactions between high and low disease activity compared to controls. B. Bar plots of all communication pathways based on number. Red are significantly more present in RA (low or high disease activity) and blue are significantly more present in controls. C. Bar plots of all communication pathways based on interaction strength.** DA, Disease Activity; RA, Rheumatoid Arthritis

**Figure S12 | A. Table of the ligand-receptor pairs that contribute to the communication for the PECAM1, HGF, IFN-II, NT and VISTA pathways. B. Bar plots showing the relative contribution of the ligand-receptor pairs that are involved in the communication for the VEGF and IL2 pathways for high and low disease activity and controls.** DA, Disease activity; L, Ligand; R, Receptor.

**Figure S13 | Heatmaps of the relative importance of cells as senders and receivers for the VEGF, IL2, HGF, and PECAM1, NT signaling pathway network in high and low disease activity. Controls are also shown for VEGF and IL2.** CD, Cluster differentiation; DC, Dendritic cells, IFIT, Interferon Induced proteins with Tetratricopeptide repeats; IFITM, interferon-induced transmembrane; Tem, T effector memory; TEMRA, Terminally differentiated effector memory.

**Table S1 | Clinical characteristics between Rheumatoid arthritis patients and matched controls.** The p-value threshold for significance was set at <0.05.Student's t-test was conducted to analyze continuous variables. For categorical variables, a Chi-square test was performed. SD, standard deviation

**Table S2 | List of differentially expressed genes between RA and matched controls and according to cell subtype. FDR ≤ 0.05, |log2(FC)| ≥ 1.6, 0.08 < base mean < 4.** CD, Cluster differentiation; DCs, Dendritics cells; IFIT, Interferon Induced proteins with Tetratricopeptide repeats; IFITM, interferon-induced transmembrane; Tem: T effector memory, TEMRA: Terminally differentiated effector memory. RA Rheumatoid Arthritis

**Figure S1 | Quality control and batch correction with Harmony.py A.UMAP projection according to sample type (control or RA) batch and lane before batch correction with Harmony. B UMAP projection according to sample type , batch and lane after batch correction with Harmony. C UMAP projection of lanes after batch correction with Harmony.** RA, Rheumatoid Arthritis; UMAP, Uniform Manifold Approximation and Projection

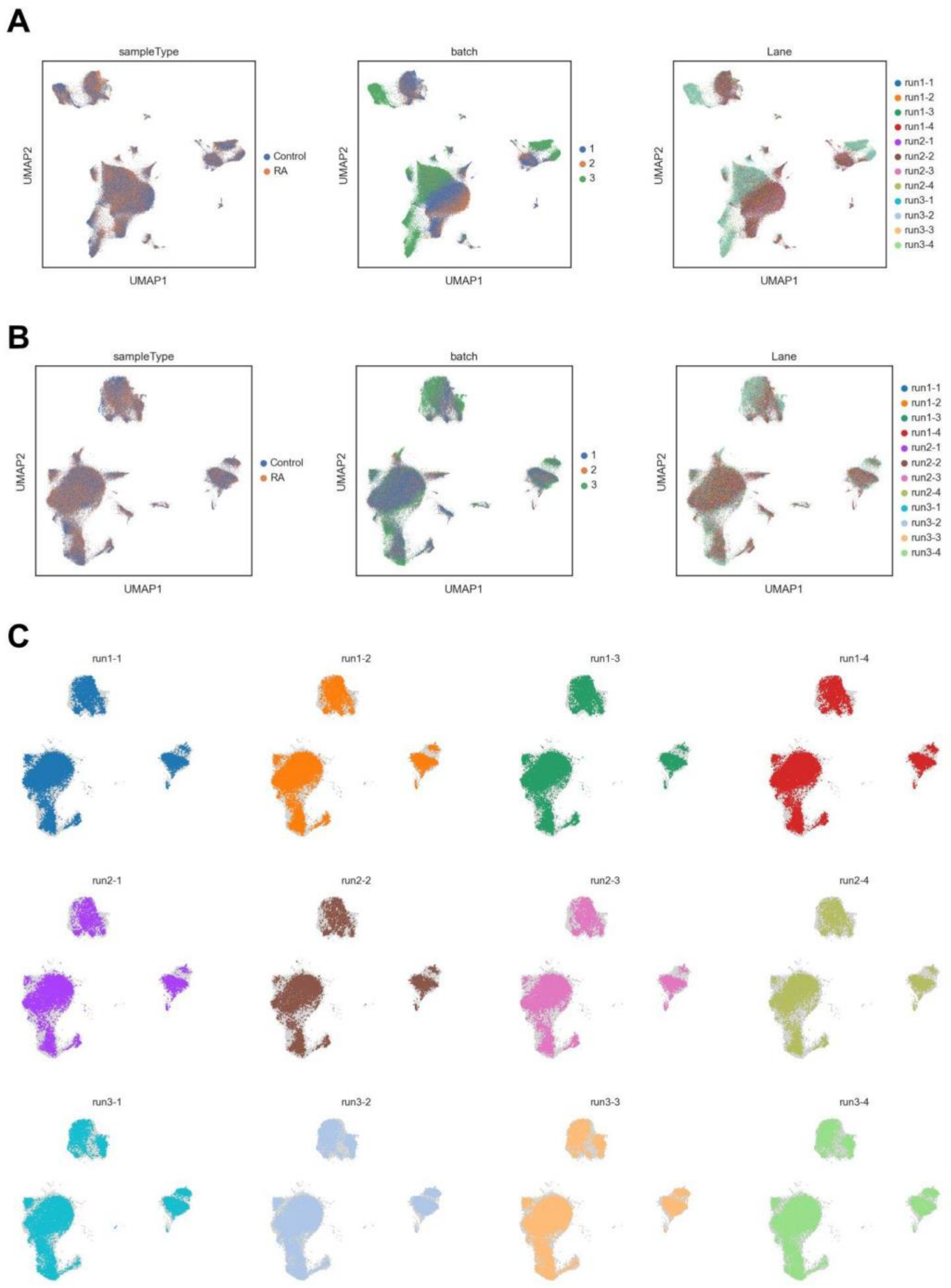

**Figure S2 | Expression heatmap representing the mean expression of B cells, CD4 T cells, CD8 T cells, NK cells and Monocytes marker genes for cell subsets.** DCs, Dendritics cells; Tem, T effector memory; TEMRA, Terminally differentiated effector memory

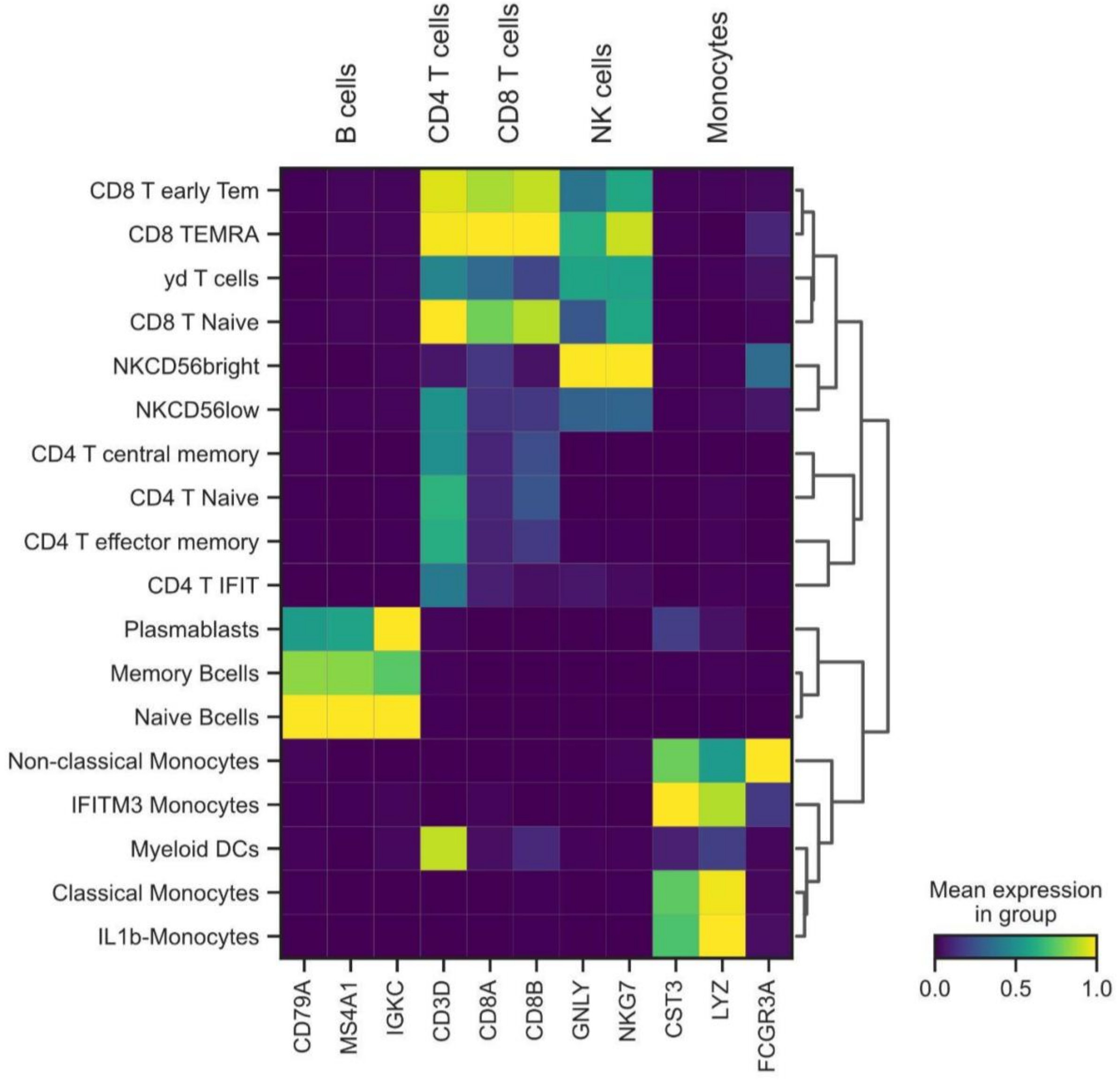

**Figure S3 | Cell proportions of B cells, CD4 T cells, CD8 T cells, NK cells and Monocytes across samples.** CD, Cluster Differentiation; NK, Natural Killer; RA, Rheumatoid Arthritis

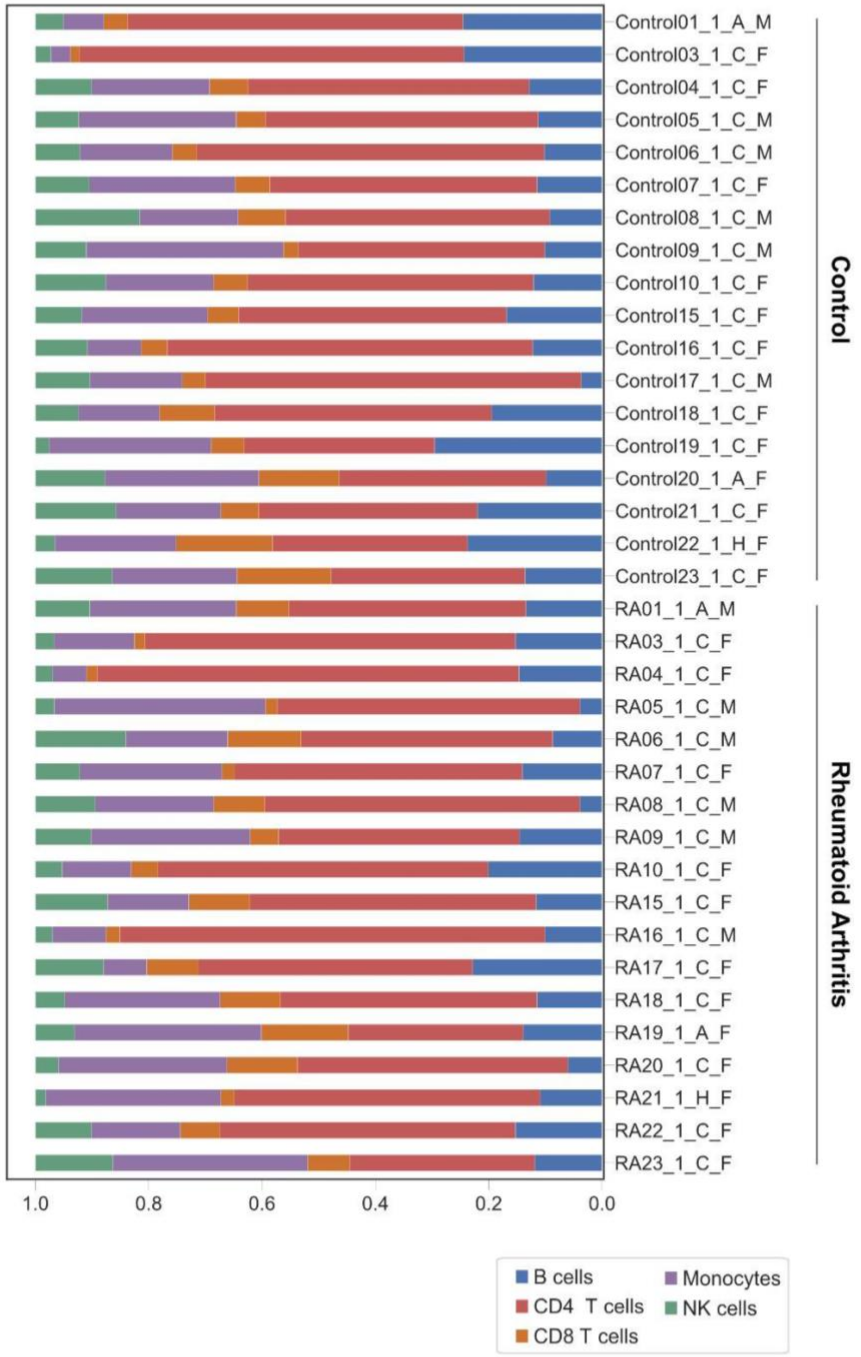

**Figure S4 | A. UMAP representation of IFITM3 gene expression across all PBMCs cell types. B. UMAP visualization of IFITM3 gene expression across monocyte subsets.** CD: Cluster Differentiation; DCs: Dendritic Cells; IFIT: Interferon Induced proteins with Tetratricopeptide repeats; IFITM: Interferon-induced Transmembrane proteins; Tem: T Effector Memory; TEMRA: Terminally Differentiated Effector Memory. RA: Rheumatoid Arthritis

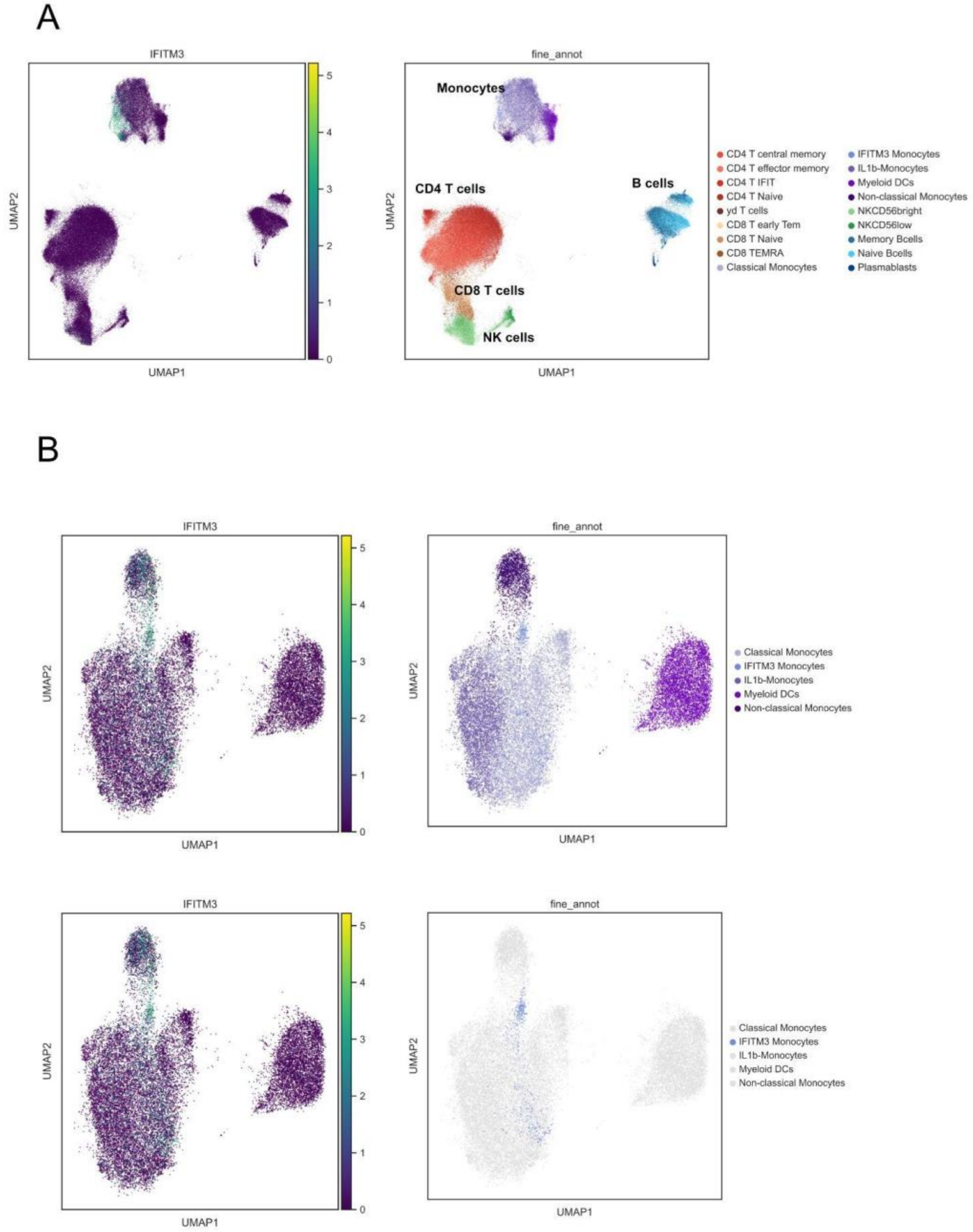

**Figure S5 | A. Cell subsets for B cells, Monocytes, CD4 T cells, CD8 T cells, NK cells. B. Compositional and density analysis between patients with Rheumatoid Arthritis and matched controls. C. Cell proportion analysis between patients with Rheumatoid Arthritis and Controls (Mann-Whitney - Wilcoxon p≤ 0.05).** CD, Cluster differentiation; DCs, Dendritics cells; IFIT, Interferon Induced proteins with Tetratricopeptide repeats; IFITM, interferon-induced transmembrane; Tem, T effector memory; TEMRA, Terminally differentiated effector memory; RA, Rheumatoid Arthritis

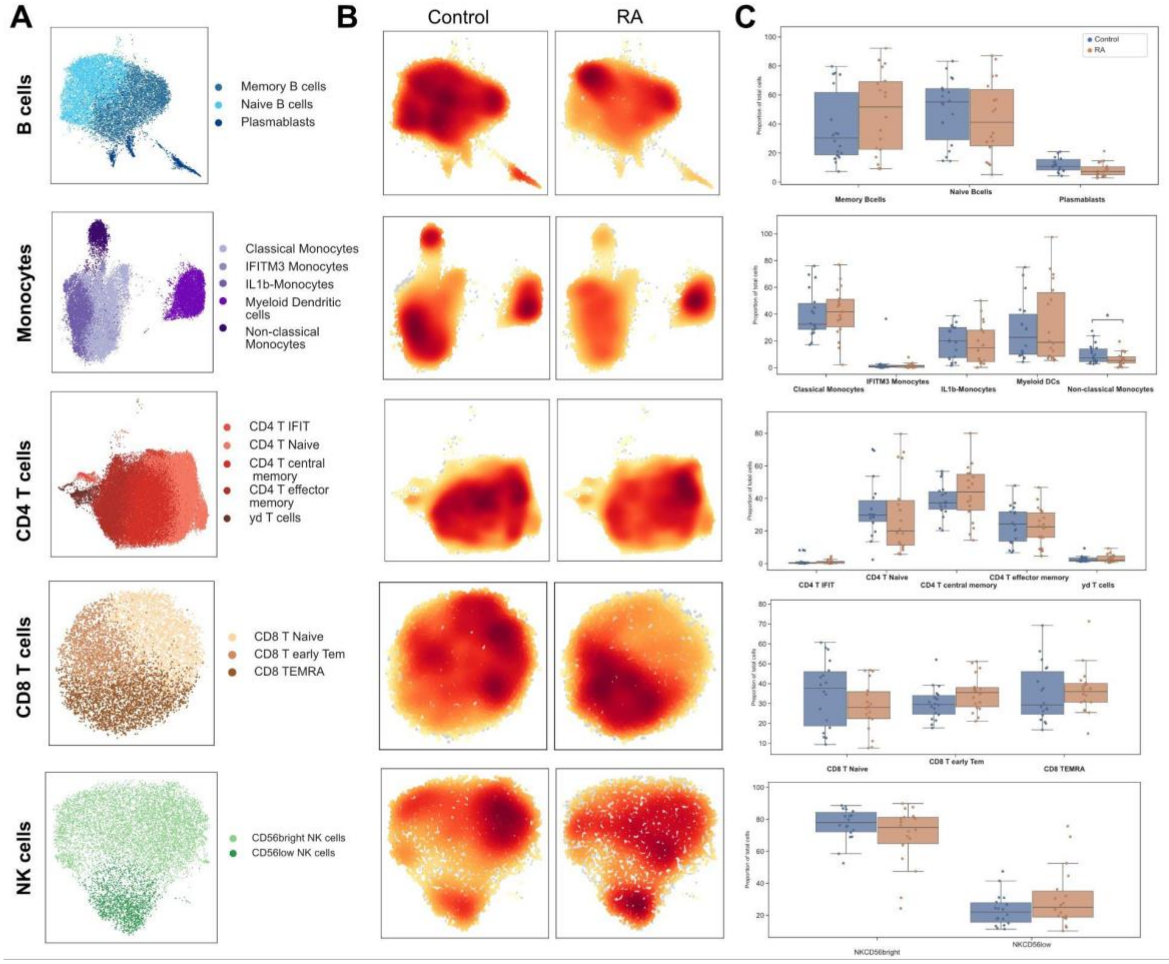

**Figure S6 | Volcano Plot representing the differential gene expression analysis between patients with Rheumatoid Arthritis and matched controls in each cell subset (FDR ≤ 0.05, log2(FC) ≥ log2(1.6), 0.08 ≤ mean expression< 4 ).** CD, Cluster differentiation; DCs, Dendritic cells; FC, Fold change; FDR, False discovery rate; IFIT, Interferon Induced proteins with Tetratricopeptide repeats; IFITM, interferon-induced transmembrane; Tem, T effector memory; TEMRA, Terminally differentiated effector memory; RA, Rheumatoid Arthritis

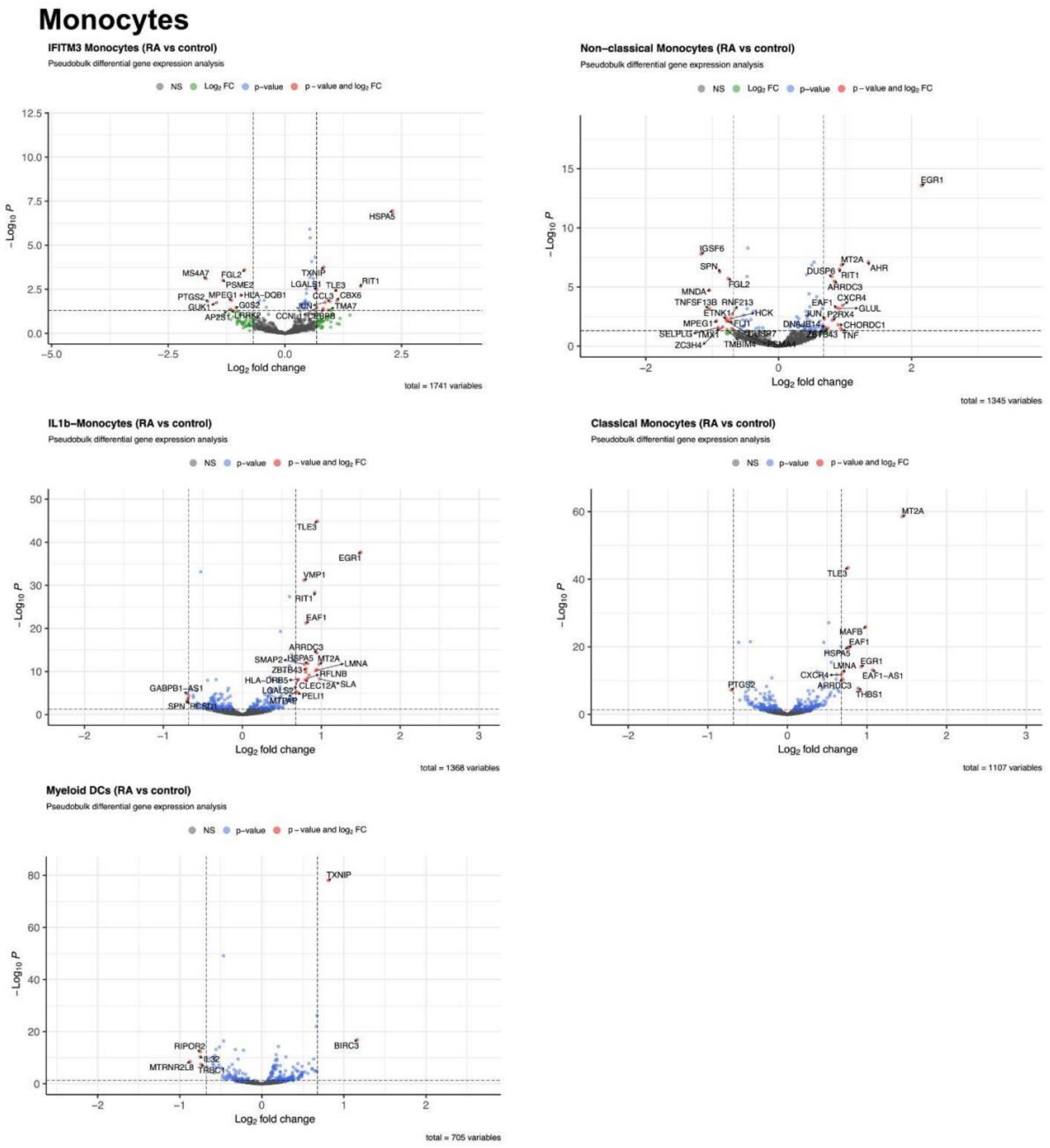

### CD4 T cells

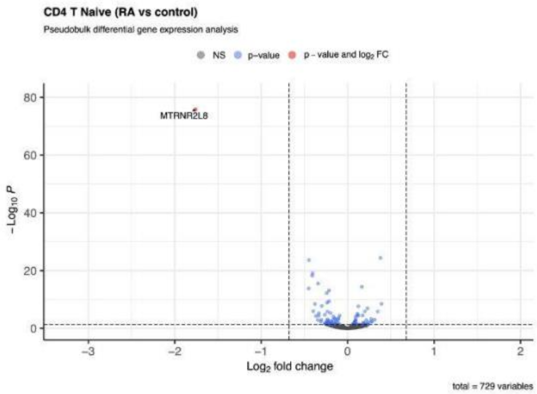


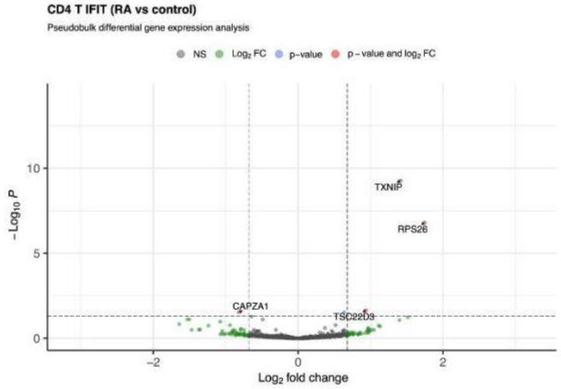


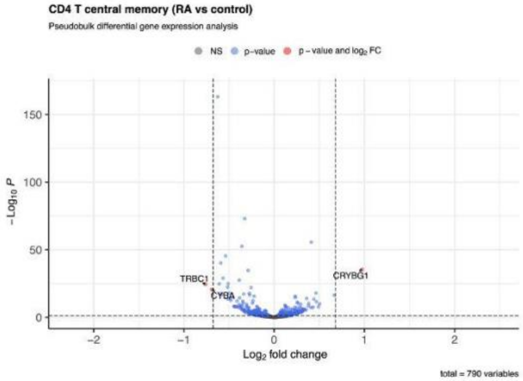


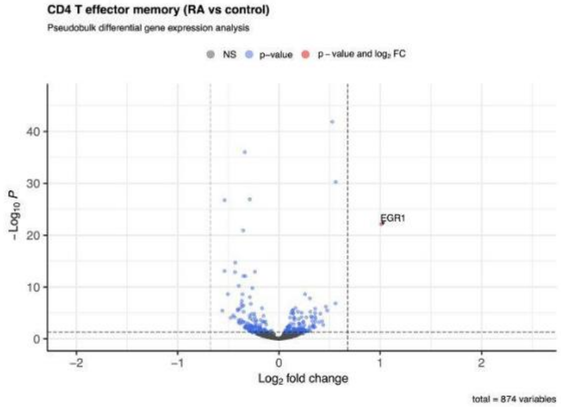


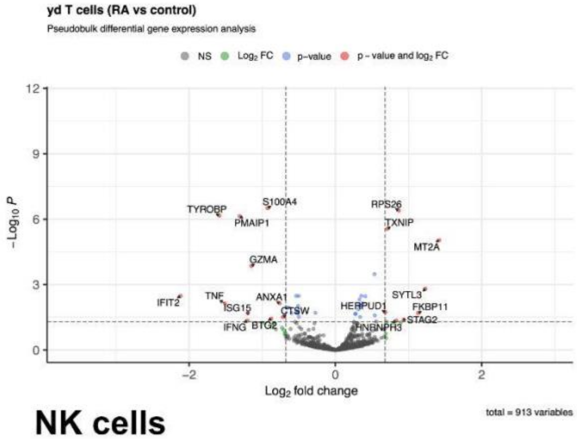


### NK cells

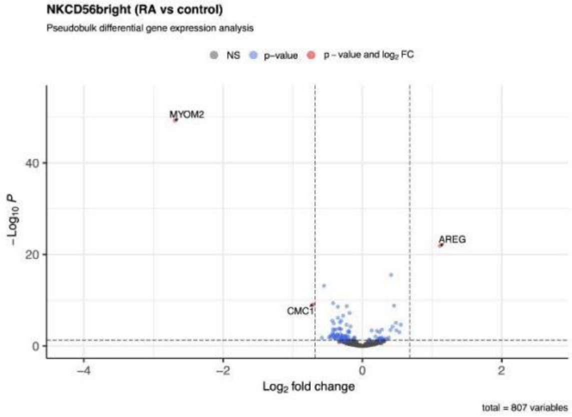


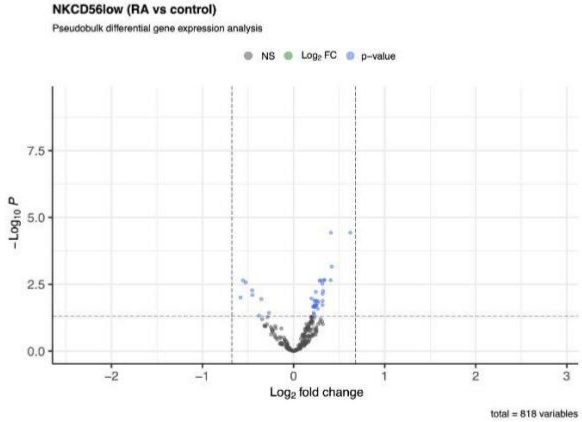

### CD8 T cells

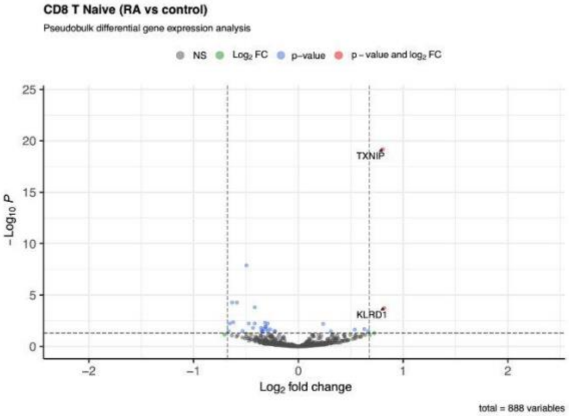


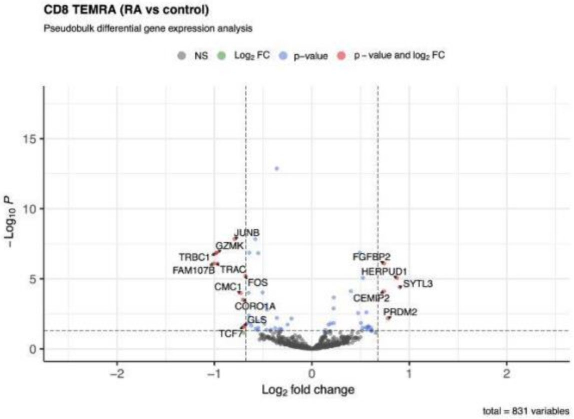


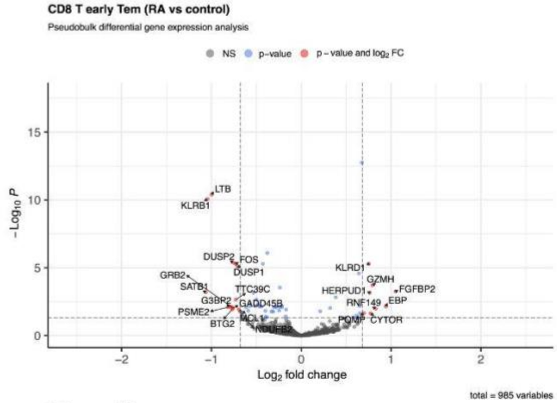


### B cells

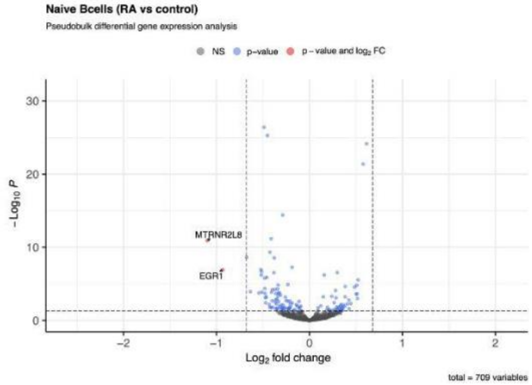


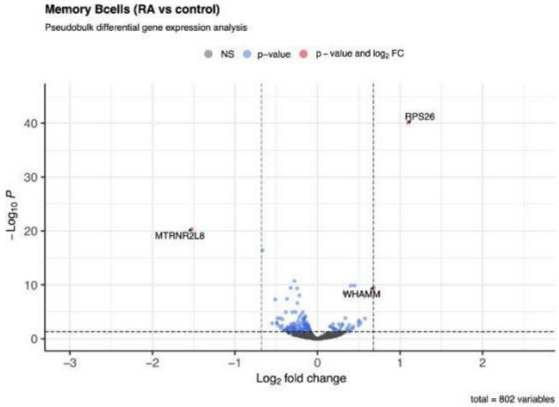


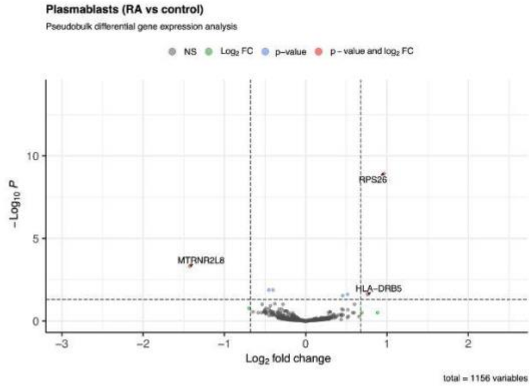

**Figure S7 | A. Spearman correlation of cell subset proportion with the DAS28-CRP for each subset. B. Correlation heatmap of cell subset and DAS-28-CRP.** CD, Cluster differentiation; DC, Dendritic cells' IFIT, Interferon Induced proteins with Tetratricopeptide repeats; IFITM, interferon-induced transmembrane; Tem, T effector memory; TEMRA, Terminally differentiated effector memory.

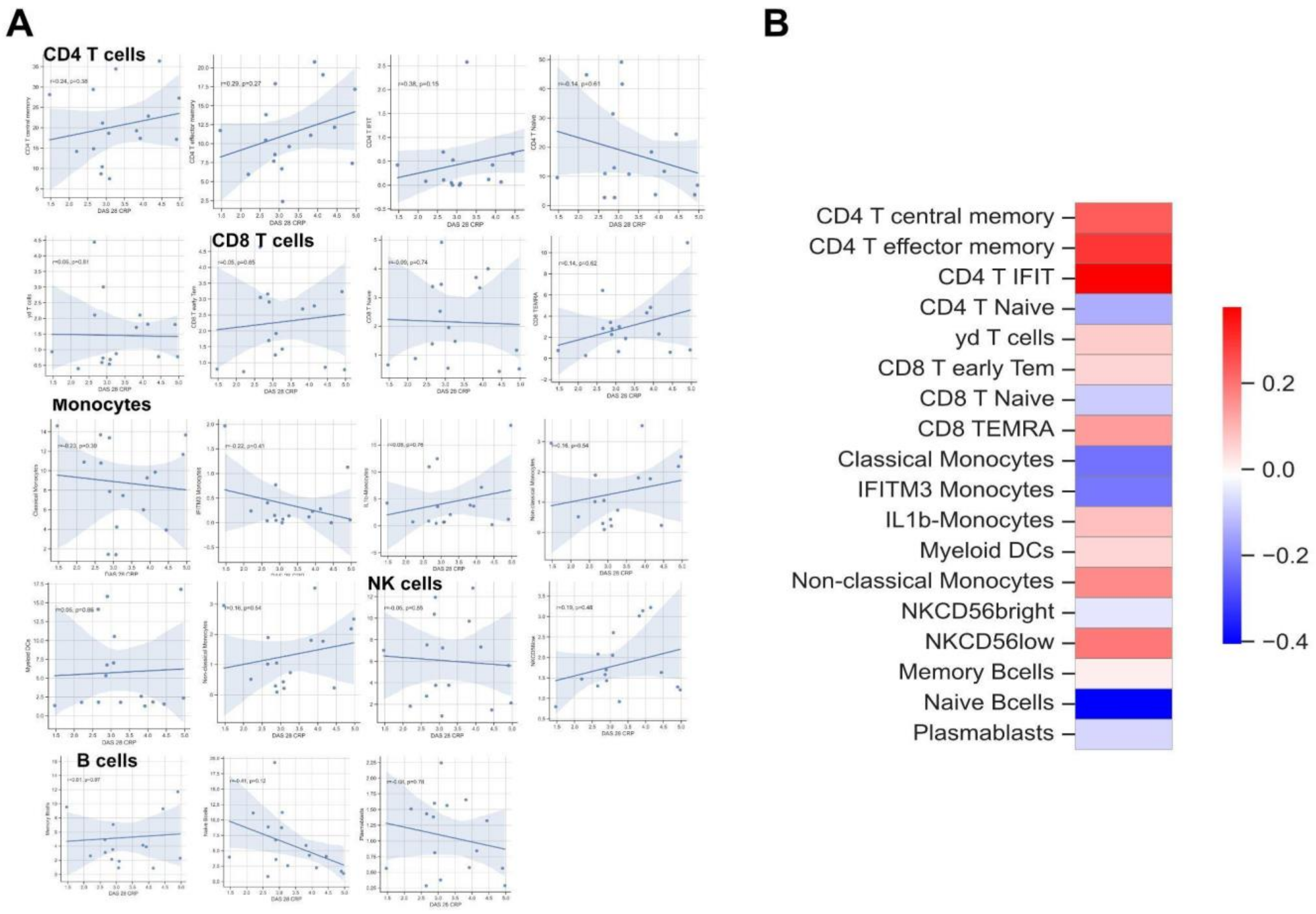

**Figure S8 | Venn Diagram of the genes differentially expressed in RA low and high disease activity compared to controls. (FDR ≤ 0.05, log2(FC) ≥ log2(1.6), 0.08 ≤ mean expression< 4 ).** RA, Rheumatoid Arthritis

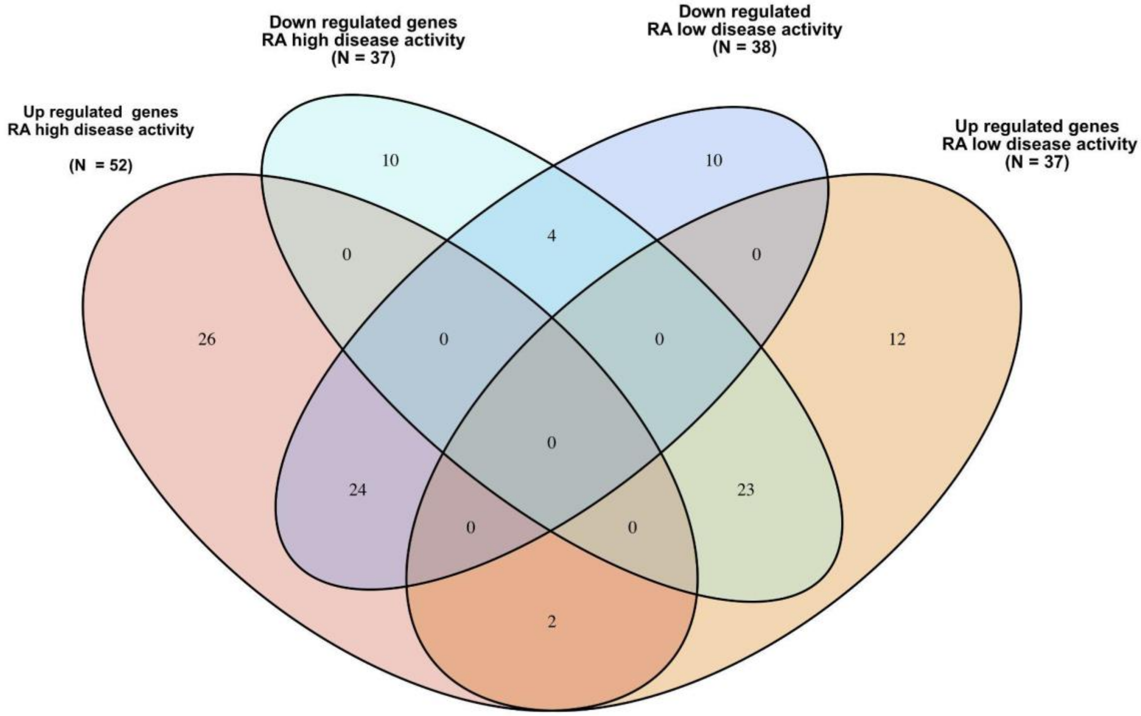

**Figure S9 | Cell-cell communications between patients with low and high disease activity and matched controls. A. Heatmap representing the relative number of interactions between RA patients with high disease activity compared to controls B. Heatmap representing the relative number of interactions between RA patients with low disease activity compared to controls.** DA Disease activity

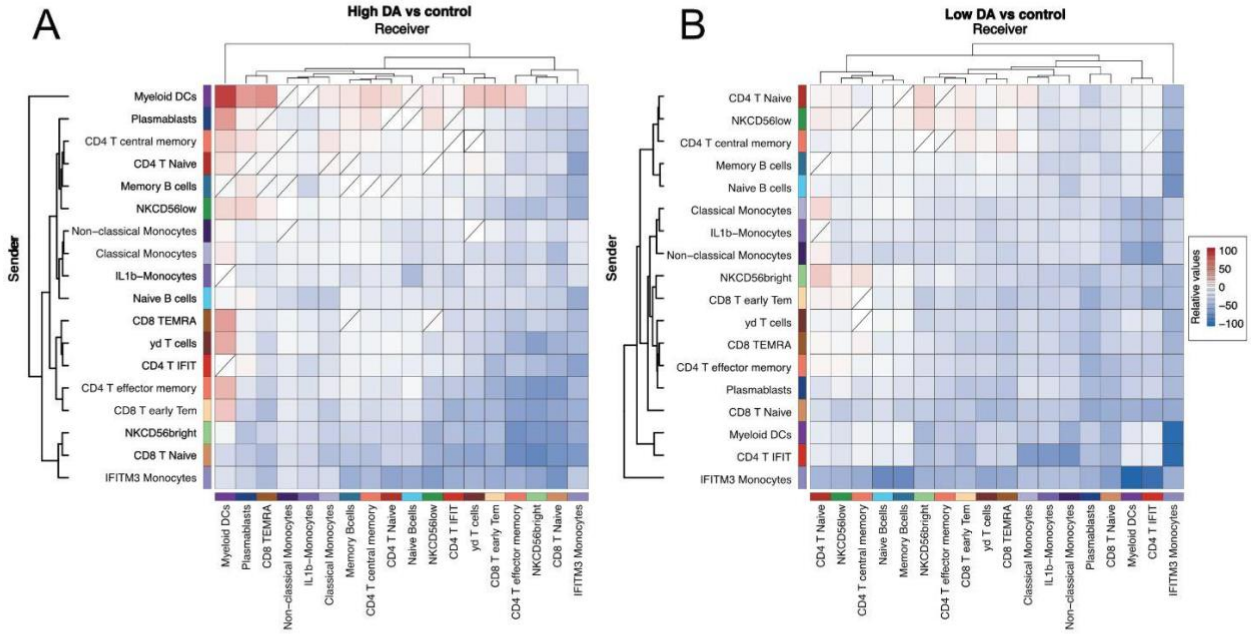

**Figure S10 | A. Bar plots of all communication pathways based on interaction strength. Red are significantly more present in RA and blue are significantly more present in controls. B. Bar plots of all communication pathways based on number.** DA, Disease Activity; RA, Rheumatoid Arthritis

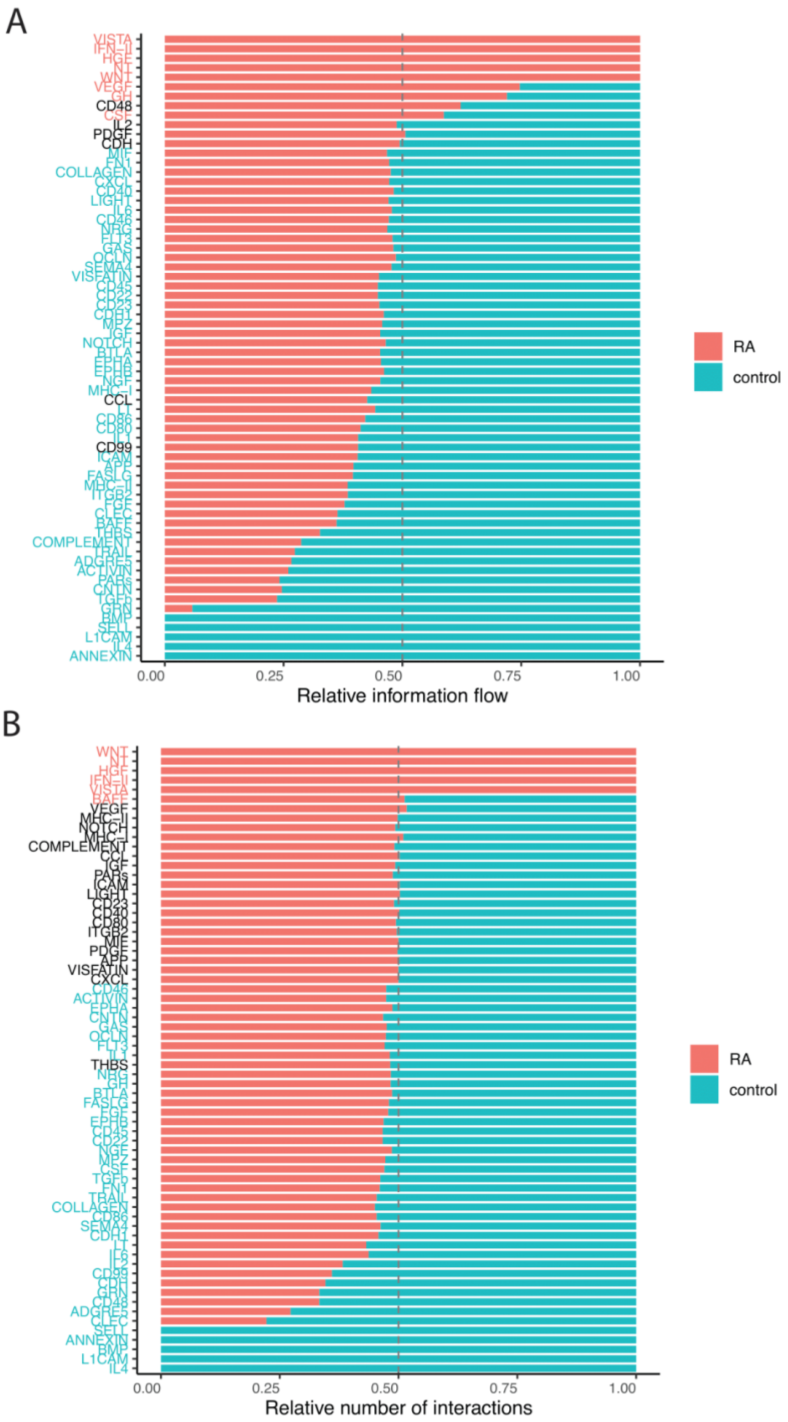

**Figure S11 | A. Dot-plot of the relative contribution of communication pathways based on number of interactions between high and low disease activity compared to controls. B. Bar plots of all communication pathways based on number. Red are significantly more present in RA (low or high Disease activity) and blue are significantly more present in controls. C. Bar plots of all communication pathways based on interaction strength.** DA, Disease Activity; RA, Rheumatoid Arthritis

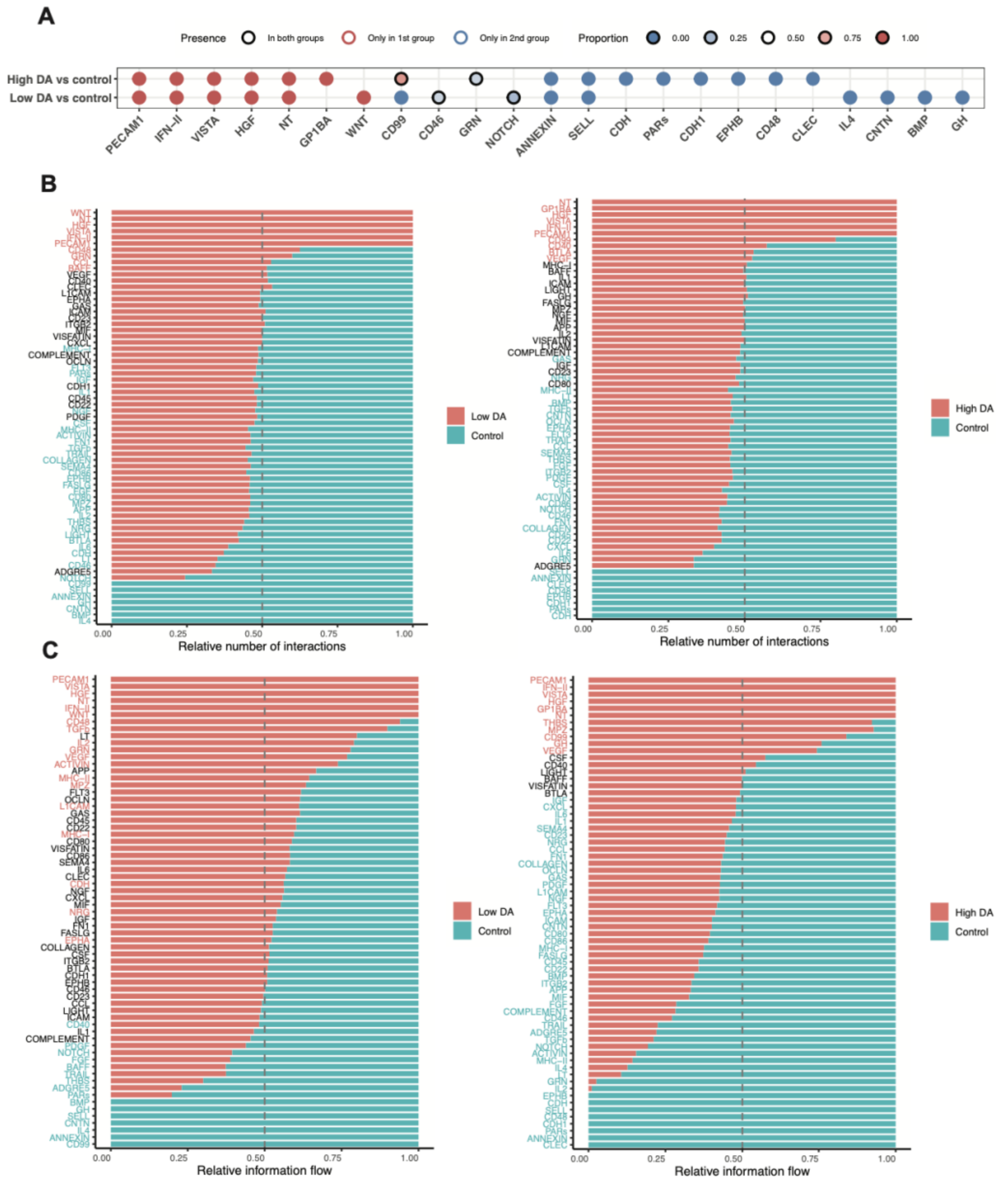

**Figure S12 | A. Table of the ligand-receptor pairs that contribute to the communication for the PECAM1, HGF, IFN-II, NT and VISTA pathways. B. Bar plots showing the relative contribution of the ligand-receptor pairs that contribute to the communication for the VEGF and IL2 pathways for high and low disease activity and controls.** DA, Disease activity; L, Ligand; R, Receptor.

**A**

| Pathway | Ligand | Receptor |
|---|---|---|
| PECAM1 | PECAM-1 | PECAM1 |
| HGF | HGF | MET |
| IFN-II | IFNG | IFNGR1+IFNGR2 |
| NT | KLK3 | NTRK1 |
| VISTA | VSIR | IGSF11 |

**B**

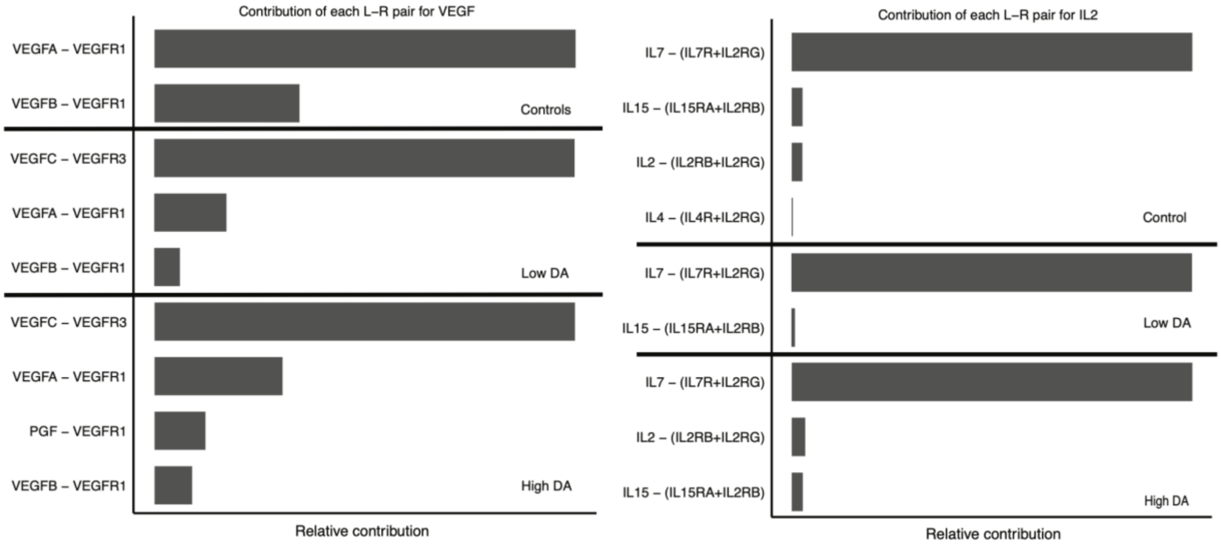

**Figure S13 | Heatmaps of the relative importance of cells as senders and receivers for the VEGF, IL2, HGF, and PECAM1, NT signaling pathway network in high and low disease activity, and for VEGF and IL2 also controls.** CD, Cluster differentiation; DC, :Dendritics cells, IFIT, Interferon Induced proteins with Tetratricopeptide repeats; IFITM, interferon-induced transmembrane; Tem, T effector memory; TEMRA, Terminally differentiated effector memory.

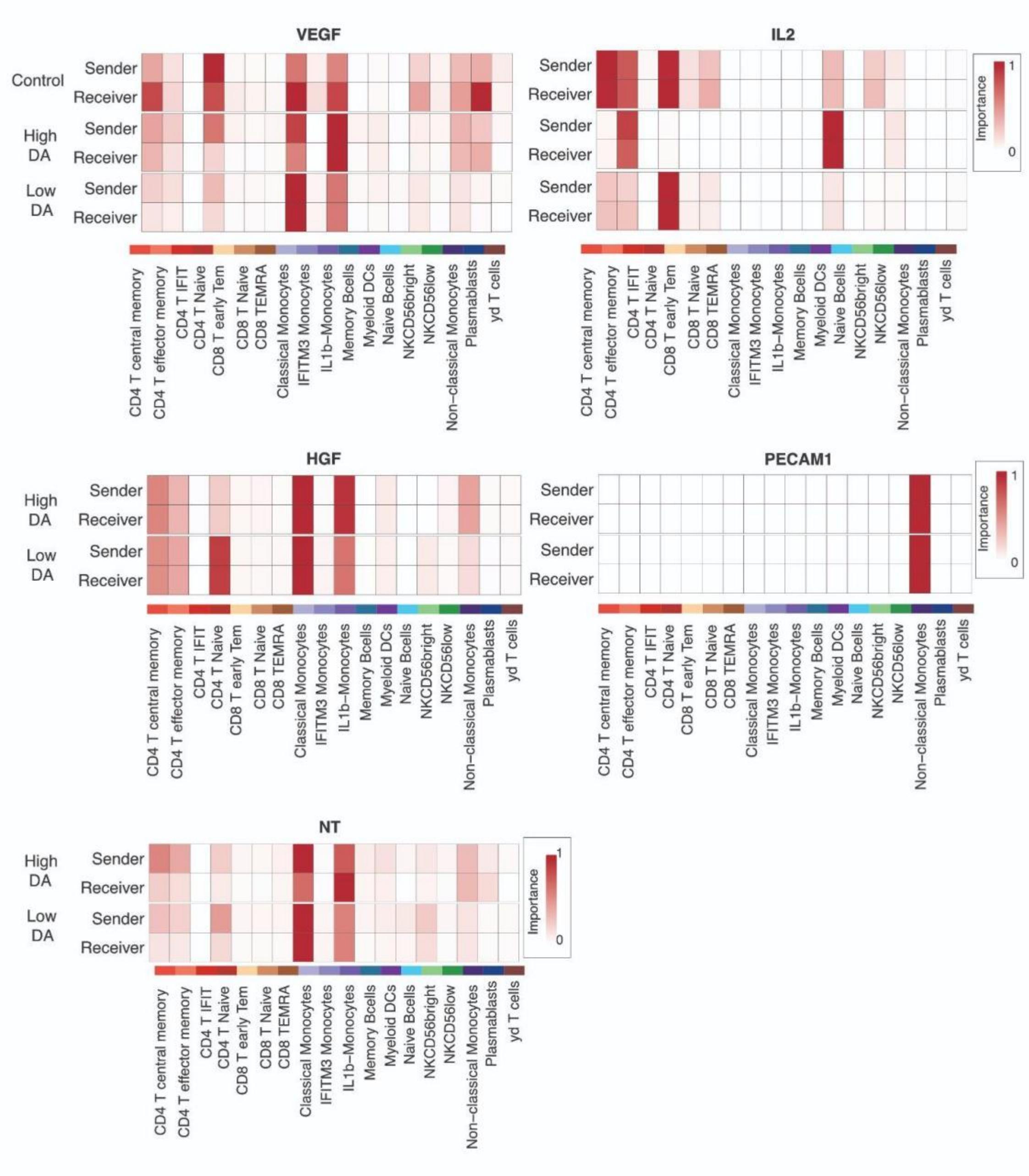

**Table S1 | Clinical characteristics between Rheumatoid arthritis patients and matched controls.** The p-value threshold for significance was set at <0.05.Student's t-test was conducted to analyze continuous variables. For categorical variables, a Chi-square test was performed. SD, standard deviation

| | | Controls (N = 18) | Rheumatoid arthritis (N = 18) | p-value | missing values (%) |
|---|---|---|---|---|---|
| **Sex** (%) | Female | 12 ( 66.7 %) | 12 ( 66.7 %) | ns | 0 |
| | Male | 6 ( 33.33) | 6 ( 33.33) | | |
| **Age (years)** (mean (SD)) | | 55.8 (16.8) | 51.7 (15.3) | ns | 11.1 |
| **Race** (%) | Asian | 2 ( 11.1 %) | 2 ( 11.1 %) | ns | 0 |
| | Caucasian | 15 ( 83.3 %) | 15 ( 83.3 %) | | |
| | Other | 1 ( 5.6 %) | 1 ( 5.6 %) | | |
| **Ethnicity** (%) | Hispanic | 1 ( 5.6 %) | 1 ( 5.6 %) | ns | 0 |
| | Not Hispanic | 17 (94.4 %) | 17 (94.4 %) | | |

**Table S2 | List of differentially expressed genes between RA and matched controls and according to cell subtype. FDR ≤ 0.05, |log2(FC)| ≥ 1.6, 0.08 < base mean < 4.** CD, Cluster differentiation; DCs, Dendritics cells; IFIT, Interferon Induced proteins with Tetratricopeptide repeats; IFITM, interferon-induced transmembrane; Tem: T effector memory, TEMRA: Terminally differentiated effector memory. RA Rheumatoid Arthritis

| Cell type | Cell subtype | Upregulated genes | Downregulated genes |
|---|---|---|---|
| **B cells** | Memory B cells | 1 | 1 |
| | Naive B cells | 0 | 2 |
| | Plasmablasts | 1 | 1 |
| **Monocytes** | Classical Monocytes | 11 | 1 |
| | IFITM3 Monocytes | 11 | 1 |
| | IL-1β Monocytes | 21 | 3 |
| | Myeloids DCs | 2 | 4 |
| | Non-classical Monocytes | 15 | 16 |
| **CD4 T cells** | CD4 IFIT | 2 | 1 |
| | CD4 Naive T cells | 0 | 1 |
| | CD4 T central memory | 1 | 2 |
| | CD4 T effector memory | 1 | 0 |
| | γδ T cells | 7 | 11 |
| **CD8 T cells** | CD8 T Naive | 2 | 0 |
| | CD8 T early Tem | 8 | 14 |
| | CD8 TEMRA | 5 | 10 |
| **NK cells** | NKCD56bright | 1 | 2 |
| | NKCD56low | 0 | 0 |
